# Evolutionary rewiring of regulatory topology refines butterfly wing patterns

**DOI:** 10.64898/2026.10.11.758279

**Authors:** Shen Tian, Yifei Joye Zhou, Anyi Mazo-Vargas

## Abstract

Phenotypes evolve through changes in gene regulatory networks (GRNs), but how changes in network topology contribute to phenotypic evolution is still poorly understood. Butterfly wing color patterns provide a tractable system to answer this question. Here, we characterized spatiotemporal expression of four color pattern genes, *WntA, pdm3, ivory*, and *Optix*, in five nymphalid butterflies, and tested their regulatory connections in WntA-associated patterns through genetic perturbations in three species spanning 80 million years of divergence. We found broadly conserved 1) regulatory hierarchy linking early pattern instruction via WntA signaling to subsequent color readout via *ivory* and *Optix*, 2) positive autoregulation in WntA signaling, and 3) positive regulation from WntA signaling to *pdm3*. The WntA-pdm3 axis, however, produced detectable color pattern readouts only in *Vanessa cardui*. In this species, additional negative feedback from *pdm3* to WntA signaling closes a reciprocal *WntA*-*pdm3* feedback loop that refines spatially separated pattern elements. We show that developmental programs can evolve by rewiring internal interactions among pre-existing GRN nodes, emphasizing the importance of network topology in phenotypic evolution.

## Introduction

Understanding how phenotypes evolve via changes in gene regulatory networks (GRNs) is a classic theme in evolutionary-developmental (evo-devo) biology. Phenotypes evolve when genes and their associated GRNs are redeployed across developmental contexts or environmental conditions, often via cis-regulatory changes^1-4^. Classic examples include the origin or loss of complex traits or environmentally responsive phenotypes, such as the emergence of butterfly eyespots, loss of snake limbs, or the origin of fly circadian plasticity^5-20^. While these examples highlight GRN redeployment as a well-established mechanism of phenotypic diversification, much less is known about how rewiring the internal regulatory interactions, or edges, across pre-existing GRN nodes can also generate diverse developmental programs and phenotypes.

A great morphological diversity and accumulating knowledge on the underlying genetic machineries make butterfly wing color patterns an ideal system to study GRN evolution. Nymphalidae is the biggest butterfly family that diversified 80 million years ago, leading to over 6000 extant species^21^. In this family, a proposed nymphalid ground plan describes diverse wing patterns as series of homologous ‘symmetry systems’ with conserved pattern elements^22^(fig. 1 left column). This morphological homology, in fact, can be explained at the molecular level. A well-studied example is that the expression of a gene encoding a Wnt-signaling ligand, *WntA*, prefigures the conserved central symmetry system (CSS), marginal band system (MBS), as well as patterns highly departed from the ground plan^23-30^(fig. 1). WntA ligand signals pattern formation via membrane receptor Frizzled 2 (Fz2) during early pupal development^31^. Disrupting *WntA* causes extensive pattern modifications across nymphalids^23,26,28,31^, and changes in its cis-regulatory modules underlie pattern variations across evolutionary scopes^25,27,32,33^. These findings suggest a flexible regulatory architecture surrounding WntA signaling that can be continuously tinkered during evolution to generate morphological diversity.

**Fig. 1.**
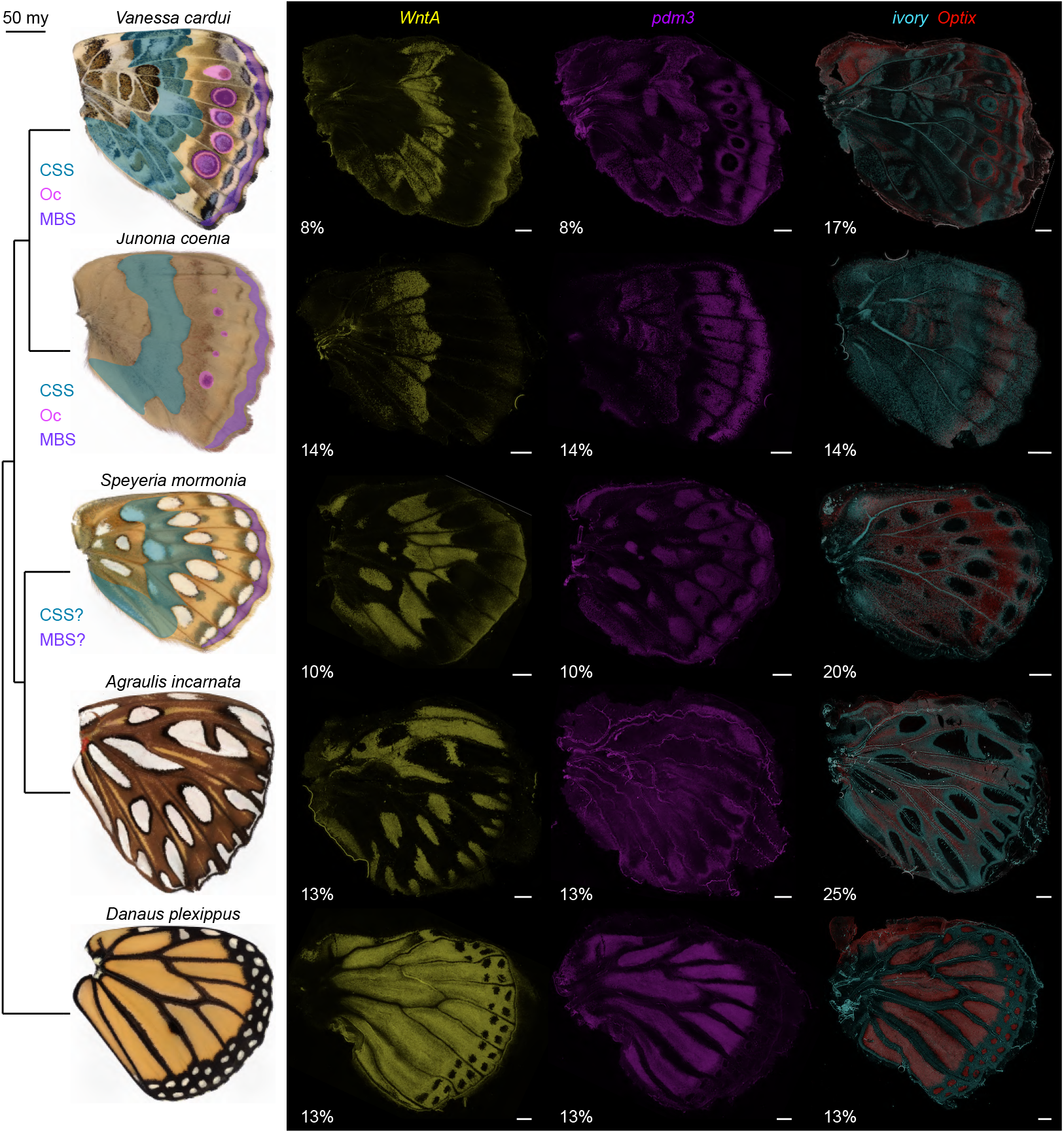
Spatial expression domains of key color pattern genes across nymphalid butterflies. Adult ventral hindwings and HCR staining of key butterfly color pattern genes, *WntA, pdm3, ivory*, and *Optix*, on the ventral surfaces of pupal hindwings across five representative nymphalid butterflies are shown. Common color pattern systems or elements described in a proposed nymphalid wing pattern ground plan are labeled. Central symmetry system (CSS), Ocelli (Oc), Marginal band system (MBS). Percentages refer to pupal developmental stages (table S2). Numbers of replicates for HCR staining are summarized in table S3. Scale bar, 500 microns.

Several candidate genes are potentially involved in the WntA-associated GRN. One of such candidates is *pdm3*, encoding a POU-domain transcription factor associated with cuticle color variation in *Drosophila* flies and beetles^34,35^, whereas its role in butterfly wing color pattern has only recently emerge. In *Vanessa cardui* and *Junonia coenia, pdm3* is expressed across multiple wing pattern elements, and its disruption causes extensive changes in colors and patterns, including the WntA-associated CSS in *V. cardui*^36,37^. This suggests a potential regulatory link across *pdm3* and WntA signaling. Beyond *pdm3*, WntA signaling may coordinate scale-color specification through *ivory* and *Optix*, conserved master regulators of melanin (black or dark) and ommochrome (red or orange) pigmentation, respectively^38-40^. In *Heliconius*, WntA signaling restricts the spatial domain of *Optix* expression^28^, but the broader regulatory relationships linking WntA signaling to these color regulators remain unresolved.

A flexible regulatory architecture with promising candidate gene nodes makes WntA-associated GRN an ideal system to investigate how regulatory interactions evolve. Here, we characterized the spatiotemporal expression dynamics of *WntA, pdm3, ivory*, and *Optix* across five deeply diverged nymphalid butterflies, tested their regulatory connections within WntA-associated GRN via genetic perturbation in three species with diverse color patterns, and discussed how evolutionary changes in GRN topology fueled phenotypic evolution.

## Results

### Diverse spatial domains but conserved temporal dynamics of color pattern genes across nymphalid butterflies

Using hybridization chain reaction (HCR), we characterized the spatiotemporal expression dynamics of *WntA, fz2, pdm3, ivory*, and *Optix*, across five nymphalid species spanning approximately 80 million years of divergence: *V. cardui, J. coenia, Speyeria mormonia, Agraulis incarnata*, and *Danaus plexippus*, at 2-3 developmental timepoints across ∼6-30% pupal development (table S1-3), focusing on ventral hindwing patterns.

Our data revealed both conserved and derived spatial expression domains of *WntA* (fig. 1, S1-5). Consistent with previous studies, *WntA* expression mapped to CSS and MBS in *V. cardui* and *J. coenia*, whose wing patterns follow the nymphalid ground plan^23,24,31^. In *V. cardui*, we also detected previously unreported expression in the hindwing eyespot centers at 8% pupal development (fig. 1, S1-2). In *S. mormonia*, although ground-plan assignments are less apparent to infer based on adult phenotypes, *WntA* marked a clear medial territory consist of proximal silver spots and the adjacent interspot regions, comparable in position and shape to the CSS in *V. cardui* and *J. coenia* (fig. 1, S3). In contrast, *A. incarnata* and *D. plexippus* exhibit wing patterns highly departed from the nymphalid ground plan. As previously reported, in *A. incarnata, WntA* is expressed inside the posterior and marginal silver spots, but outside of the anterior silver spots^23,24,31^(fig. 1, S4). In *D. plexippus, WntA* is broadly expressed across the wing, with weaker expression around veins and invisible expression in the marginal white spots^23^(fig. 1, S5). Overall, our data is consistent with the reported evolutionary shifts in *WntA* expression and function^23,25^, and emphasizes how *WntA* expression could serve a developmental marker to infer molecular homology when morphological homology is hard to recognize.

Across all five species, *fz2* expression primarily complements *WntA* expression (fig. S1-5), consistent with the reported negative-feedback mechanism in which WntA signaling locally represses *fz2* transcription^31^. *pdm3* showed pattern-specific expression within each species, overlapping with *WntA* expression across several pattern elements (fig. 1, S1-5) including the CSS and MBS. These shared domains might suggest conserved but context-dependent interactions between WntA signaling and *pdm3*^36,37^. Finally, *ivory* and *Optix* were expressed across pattern elements where melanin and ommochrome pigments are presumably synthesized, respectively (fig. 1, S1-5), supporting their conserved roles as scale-color fate master regulators^38-42^.

Despite the diverse spatial domains of these genes, we observed a conserved temporal dynamic in expression – *WntA, fz2*, and *pdm3* are expressed earlier, by ∼6–8% of pupal development and became barely detectable after 20% pupal development. By contrast, the expression of *ivory* and *Optix* was observed simultaneously or after the activation of the three patterning genes, primarily post 10% pupal development. In some cases, *ivory* expression persisted or even intensified post 20% pupal development (fig. 1, S1-5). This temporal dynamic suggests a potential regulatory hierarchy, where WntA signaling and *pdm3* first stablish spatial boundaries of pattern elements, and *ivory* and *Optix* subsequently specify scale-color identities.

### A reciprocal feedback loop between *WntA* and *pdm3* underlies complex pattern formation in *V. cardui*

To directly draw regulatory connections among these genes, we generated mosaic knockouts (mKOs) for each target gene via embryonic CRISPR-Cas9 mutagenesis in each of *V. cardui, A. incarnata*, and *D. plexippus* (fig. S6, table S4), then examined changes in expression via in situ HCR and/or qPCR, to measure how each perturbation altered the expression of the gene disrupted and other network components (table S3).

We first focused on the ventral hindwing patterns in *V. cardui*. In this species, *WntA* and *pdm3* are co-expressed in the CSS and MBS. Within the CSS, their expression prefigures arrays of petaloid melanic bands, with an overall spatial expansion from 8% to 17% pupal development, with the later expression closely resembling the adult patterns organization (fig. 2A, S1). *pdm3* is additionally expressed in several *WntA*-negative elements including eyespot rings and discal elements, as reported previously^36^ (fig. 2A, S1). Here, we focus on the CSS, a system where WntA is the established patterning signal.

**Fig. 2.**
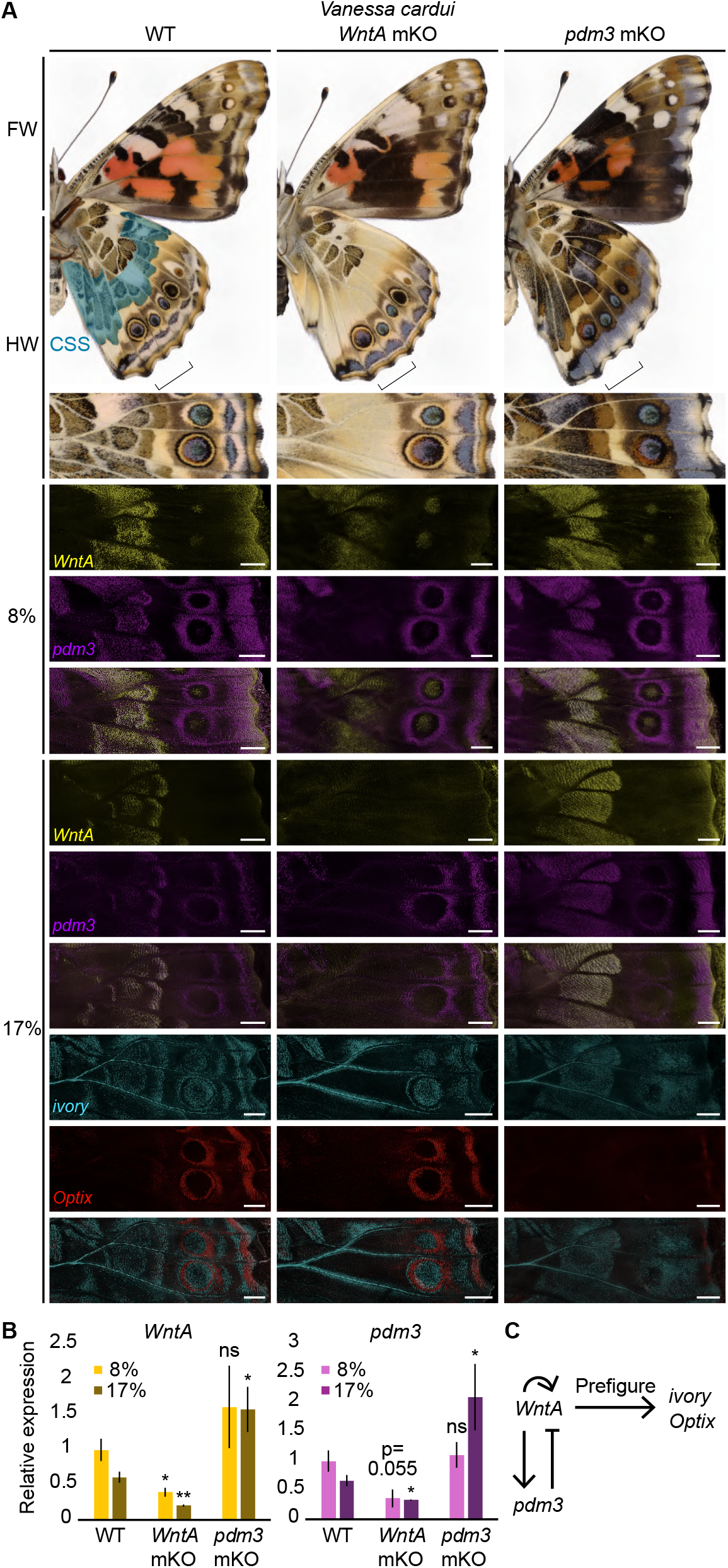
A reciprocal feedback loop across *WntA* and *pdm3* underlies complex ventral hindwing CSS pattern formation in *V. cardui*. **(A)** Adult ventral surfaces and HCR staining on the ventral surfaces of pupal hindwings are shown across WT, *WntA* mKO, and *pdm3* mKO in *V. cardui*. CSS, central symmetry system. Numbers of replicates for HCR staining are summarized in table S3. Scale bar, 500 microns. **(B)** HCR-stained whole hindwings were genotyped via Sanger sequencing and those with high knockout rates were used for qPCR (table S5). Relative expression levels of *WntA* and *pdm3* were measured across WT, *WntA* mKO, and *pdm3* mKO (n= 6 (8% WT), 3 (17% WT), 3 (8% *WntA* mKO), 3 (17% *WntA* mKO), 4 (8% *pdm3* mKO), 4 (17% *pdm3* mKO)). A two-tailed Student’s test was used to assess expression differences across mKO and WT in each stage. ns, not significant; *p < 0.05; **p < 0.01; ***p < 0.001. Error bar, SEM. **(C)** Wing color pattern GRN was proposed in *V. cardui*. In **(A)** and **(B)**, percentages refer to pupal developmental stages.

Disrupting *WntA* eliminated the petaloid melanic bands in adult wings (fig. 2A). *WntA* expression was visibly weaker with less refined boundaries and internal organization at 8% pupal development, and became undetectable at 17%, supporting a previously described positive autoregulation in WntA signaling^31^ (fig. 2A, 2C, S7). Strikingly, *pdm3* expression was completely depleted in the CSS, suggesting that WntA signaling is required to induce *pdm3* expression (fig. 2A, 2C, S7). Conversely, *pdm3* disruption caused the petaloid bands to fuse, eliminated their internal spacing and increased melanization (fig. 2A). Aligning with this phenotype, *WntA* and *pdm3* expression fused, becoming homogeneous across the CSS (8% and 17%), with greater overall signal intensity at 17% (fig. 2A, S8). These results indicate that Pdm3 represses WntA signaling to generate the refined CSS pattern details (fig. 2C).

To quantitively validate these interactions, we co-extracted genomic DNA and total RNA from HCR-stained mKO wings, genotyped each sample and used those with high mutation rates to quantify gene expression via qPCR (table S5-6). Although the whole wing measurements may underestimate spatially restricted effects, the data supported the in-situ observations: *WntA* disruption significantly (p<0.05, two-tailed Student’s test) reduced the expression of *WntA* (8% and 17%) and *pdm3* (8% (p=0.055) and 17%), while loss of *pdm3* significantly increased *WntA* and *pdm3* expression at 17% development (fig. 2B)

We then tested how *WntA* and *pdm3* impacts the expression of *ivory* and *Optix*. In both *WntA* and *pdm3* mKO wings, we observed correlated expression changes in *ivory* and *Optix* with the corresponding adult phenotypes, suggesting that WntA signaling and *pdm3* instruct scale color fates via these color regulators (fig. 2A, S7-8). In contrast, disrupting *ivory* did not produce visible expression changes in *WntA* or *pdm3*, suggesting minimal feedback from the color regulators, at least for *ivory*, to upstream patterning machineries (Supplementary text 1, fig. S9. table S7).

Overall, we discovered in *V. cardui* a reciprocal *WntA*-*pdm3* feedback loop that prefigures the internal complexity of the CSS and orchestrates downstream scale color regulators, *ivory* and *Optix* (fig. 2C).

### The *WntA* – *pdm3* feedback loop is not closed in *A. incarnata* and *D. plexippus*

To test whether the *WntA*-*pdm3* feedback loop is evolutionarily conserved, we examined two other nymphalids with wing patterns highly departed from the nymphalid ground plan, *A. incarnata* and *D. plexippus*.

In *A. incarnata, WntA* and *pdm3* are co-expressed in the posterior silver spots, marginal silver spots, and brown regions surrounding the anterior silver spots. *pdm3* expression also extended across a broad vein-associated domain outside the WntA-positive regions (8% and 13%) (fig. 3A, S4). In *WntA* mKOs, posterior and marginal silver spots were depleted, while anterior silver spots expanded (fig. 3A). *WntA* expression was strongly reduced or almost undetectable at 13% (fig. 3A, S10), supported by qPCR (fig. 3B, table S5-6). These data support a conserved positive autoregulation in WntA signaling^31^ (fig. 3C). Meanwhile, *pdm3* expression appeared completely undetectable across all WntA-positive regions but remained visibly unchanged around the veins (fig. 3A, S10), supporting a conserved positive regulation from WntA signaling to *pdm3* (fig. 3C). Whole wing qPCR did not capture this localized reduction, likely because *pdm3* expression remained unchanged outside the WntA-positive territories (fig. 3B, table S5-6). Conversely, loss of *pdm3* produced no visibly detectable adult color patterns changes (fig. 3A, S11, table S8) and did not alter *WntA* or *pdm3* expression in-situ or in qPCR (fig. 3A-B, S12, table S5-6), providing minimal evidence for a *pdm3*-to-*WntA* feedback.

**Fig. 3.**
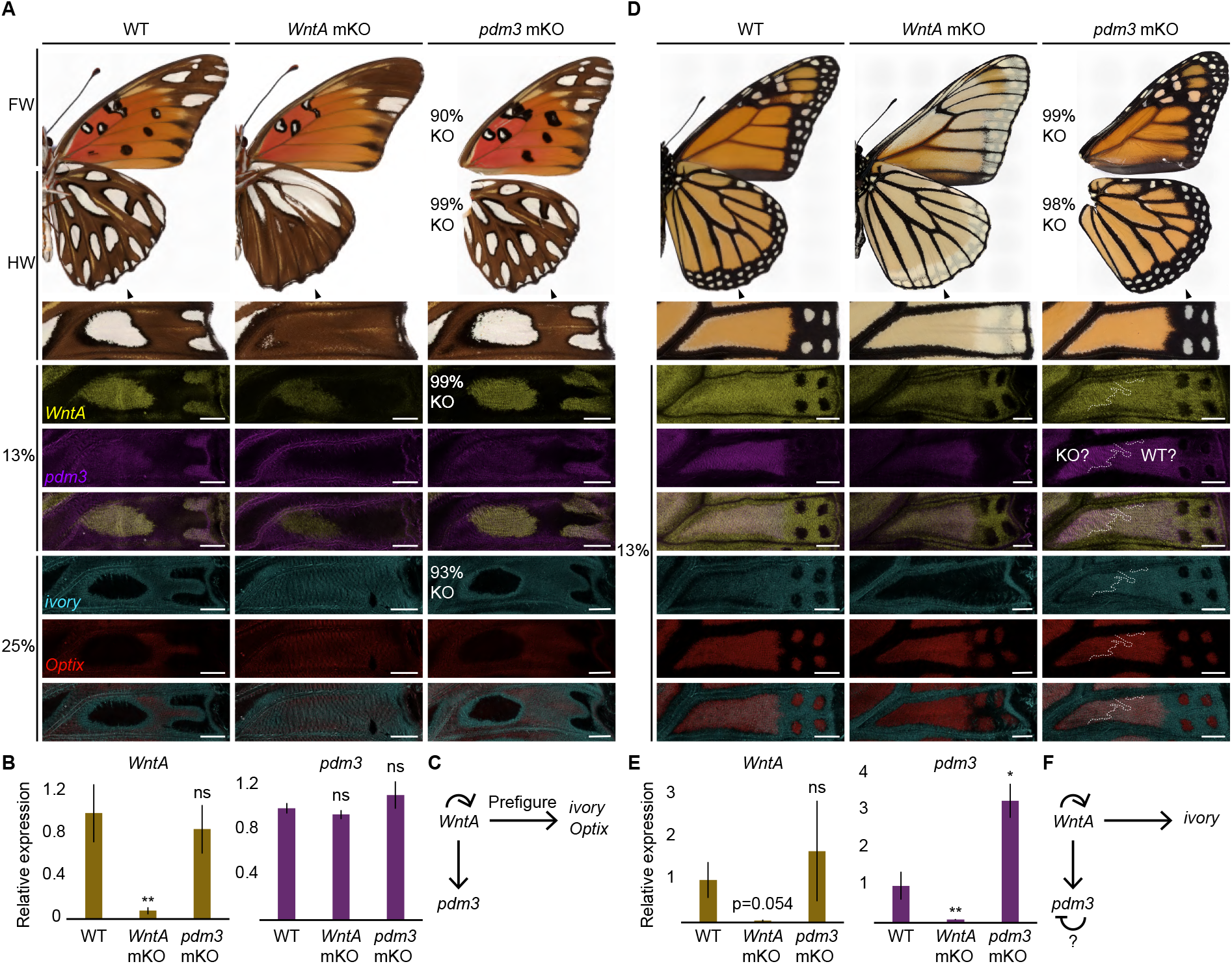
The *WntA*-*pdm3* feedback loop is not closed in *A. incarnata* and *D. plexippus*. **(A-C)** *A. incarnata* and **(D-F)** *D. plexippus* were used for comparative investigations. Adult ventral surfaces and HCR staining on the ventral surfaces of pupal hindwings are shown across WT, *WntA* mKO, and *pdm3* mKO in *A. incarnata* **(A)** and *D. plexippus* **(D).** For adult/pupal wings without visible color/expression changes, wings were genotyped via Sanger sequencing and knockout (KO) rates were shown as indel rates. Percentages refer to pupal developmental stages. Numbers of replicates for genotyped adult wings and HCR staining are summarized in table S8 and table S3, respectively. Scale bar, 500 microns. HCR-stained whole hindwings were genotyped via Sanger sequencing and those with high knockout rates were used for qPCR (table S5). Relative expression levels of *WntA* and *pdm3* were measured across WT, *WntA* mKO, and *pdm3* mKO in *A. incarnata* (n= 5 (WT), 5 (*WntA* mKO), 5 (*pdm3* mKO)) **(B)** and *D. plexippus* (n=6 (WT), 3 (*WntA* mKO), 3 (*pdm3* mKO)) **(E)**, both at 13% pupal development. A two-tailed Student’s test was used to assess expression differences across mKO and WT in each species. ns, not significant; *p < 0.05; **p < 0.01; ***p < 0.001. Error bar, SEM. Wing color pattern GRNs were proposed in *A. incarnata* **(C)** and *D. plexippus* **(F)**.

These results suggest that while a positive WntA autoregulation and a positive *WntA*-to-*pdm3* regulation remain conserved, the negative *pdm3*-to-*WntA* regulation present in *V. cardui* is not detectable in *A. incarnata* (fig. 3C).

In *D. plexippus*, a basal nymphalid lineage, *WntA* was broadly expressed across the wing except the marginal white spots, whereas *pdm3* was expressed in a subset of WntA-positive regions in the intervein orange regions (6% and 13%) (fig. 3D, S5). In *WntA* mKOs, the intervein orange regions and the marginal black bands appeared strongly washed out, while the black stripes around wing veins remained unchanged. Both *WntA* and *pdm3* showed reduced expression levels at 13% pupal development (fig. 3D, S13), supported by qPCR (p=0.054 for *WntA* in *WntA* mKO) (fig. 3E, table S5-6). As in *V. cardui* and *A. incarnata*, these results are consistent with a conserved positive autoregulation in WntA signaling, and a conserved positive regulation from WntA signaling to *pdm3* (fig. 3F). As in *A. incarnata*, loss of *pdm3* showed no visible changes in adult color patterns (fig. 3D, S14, table S8). Across *pdm3* mKOs, we observed sharp boundaries between regions with different *pdm3* expression levels, potentially corresponding to WT-KO clones (fig. 3D, S15). In parallel, qPCR detected significantly increased *pdm3* expression in the mKOs (fig 3E, table S5-6), supporting a potential negative autoregulation around *pdm3* with unknown functional relevance (fig. 3F). No expression changes in *WntA* were detected either across WT and mKO tissue boundaries in-situ, or across WT and mKO wings in qPCR (fig. 3D-E, S15, table S5-6). Thus, as in *A. incarnata*, we found minimal evidence that Pdm3 feeds back on WntA signaling in *D. plexippus* (fig. 3F).

In both *A. incarnata* and *D. plexippus, WntA* loss altered *ivory* and/or *Optix* expression in spatial correspondence with the adult phenotypes, while *pdm3* mKOs showed no detectable changes in either regulator (fig. 3A, 3D, S10, S12-13, S15, table S7). Of note, in *D. plexippus*, WntA signaling broadly promotes *ivory* across its expression domains but plays a minor role in refining *Optix* expression domains (Supplementary text 2, fig. 3D, S13). These indicate a conserved regulatory cascade from WntA signaling, independently of *pdm3*, to *ivory* and *Optix* in both species (fig. 3C, 3F). As in *V. cardui*, disrupting *ivory* did not induce visible expression changes in *WntA* and *pdm3* in either species (Supplementary text 1, fig S16-S17), suggesting minimal feedback from scale color regulators to patterning machineries across nymphalids.

To sum up, across nymphalids we identified three conserved features: 1) temporal and regulatory progression from early WntA signaling to subsequent deployment of scale color regulators, *ivory* and *Optix*, 2) positive autoregulation in WntA signaling, and 3) positive regulation from WntA signaling to *pdm3*. Negative feedback from *pdm3* to WntA signaling was only detected in *V. cardui*, which closes a lineage-specific *WntA*-*pdm3* feedback loop, underlying the development of its complex CSS pattern details.

## Discussion

In this study, we characterized expression dynamics of key butterfly wing color pattern genes and their regulatory connections across deeply diverged species. Despite divergent spatial deployment and associated functions, we discovered a conserved temporal sequence when early *WntA* and *pdm3* expression preceded *ivory* and *Optix* deployment, supporting a shared regulatory progression from pattern establishment to scale colour specification in butterfly wing colour pattern development^20,28,43-45^. This may reflect a balance between developmental constraint and evolutionary flexibility: whereas spatial deployment and certain GRNs components can evolve flexibly, key regulatory hierarchy may remain conserved, potentially constraining changes that disrupt developmental organization^46-50^.

In addition to a conserved regulatory hierarchy across color pattern GRNs, we observed interesting topological changes across *WntA* and *pdm3*. First, a unidirectional positive regulation from WntA signaling to *pdm3* appears conserved across nymphalids. *pdm3* appeared to be broadly integrated into the WntA signaling network across several or all WntA-associated patterns, but not essential to produce detectable color pattern readouts across all nymphalids. This supports that gene expression and gene function can be decoupled in evolution: modular gene expression may arise through upstream cooption without detectable functional consequence, whereas additional regulatory changes might enable its functional integration^46,51,52^.

In fact, we discovered in *V. cardui* negative feedback from *pdm3* to WntA signaling, which joins the positive autoregulation in WntA signaling and the positive regulation from WntA signaling to *pdm3*, to form a functional feedback loop. In this case, *pdm3* plays an active role in color pattern regulation, likely by linking it to the downstream coloration machineries through WntA signaling. Of note, our current phylogenetic resolution does not establish whether the negative feedback was gained in *V. cardui* or lost elsewhere. And it remains unsolved whether it was established directly, or indirectly. If built directly, it could involve evolutionary changes in *WntA* cis-regulatory modules, Pdm3 activity, or its cofactors. Future studies are expected to understand when and how this regulatory edge evolved in the phylogeny.

Importantly, this feedback loop functionally contributes to the internal pattern organization in *V. cardui*. Its regulatory topology suggests that it may function as an activator-inhibitor circuit, potentially promoting self-organization and the formation of spatially separated pattern elements^53,54^. Future spatiotemporal analysis and modeling are required to elucidate the dynamic interactions between WntA signaling and Pdm3 during complex color pattern development^55-57^.

Overall, our study highlights that phenotypic evolution need not rely solely on the redeployment of genes and their associated genetic modules. Evolutionary rewiring of regulatory interactions among pre-existing GRN nodes could serve an additional route to developmental complexity and phenotypic diversity.

## Acknowledgements

We thank Kitty Bryan, Kristen Kiki Sylvain, Robert Reed lab, Carol Boggs lab, Carolina Biological Supply, Monarch Watch, and Rose Franklin’s Perennial & Butterfly Farm for supplying living butterflies. We thank Arnauld Martin for the helpful comments. We thank Duke Greenhouse staffs Jorge Fidhel Gonzalez, Amy Eddings, and Kaliany Vazquez-Hernandez for maintaining host plants.

## Materials and Methods

### Insect husbandry

Five nymphalid butterflies, *V. cardui. J. coenia, S. mormonia, A. incarnata*, and *D. plexippus*, were reared 27°C, with a 16:8 day: night cycle and 60% relative humidity in a walk-in growth chamber. A lab stock was maintained for each species except for *S. mormonia*, which was kept for one generation only. Adult butterflies were fed Gatorade. Sources and host plants/artificial diet for each species were summarized in table S1. Total pupal development time for each species are summarized in table S2.

### CRISPR-Cas9 genome editing

Embryonic CRISPR-Cas9 knock out experiments were performed in *V. cardui, A. incarnata*, and *D. plexippus*. For protein coding genes *WntA* and *pdm3*, single guide RNAs (sgRNAs) were either adopted from previous studies^31,36^ or designed using CRISPRdirect^58^, to target the coding sequence (CDS) shared by all isoforms of each gene in each species. One sgRNA was used to target each gene except for *pdm3* in *D. plexippus*, where two sgRNAs were designed and injected independently. For the long noncoding RNA *ivory*, one sgRNA was designed on the conserved *ivory* transcription start site (TSS) in each species^39^. Guide sequences were blasted against the corresponding genomes to check for specificity. All sgRNAs were ordered from Integrated DNA Technologies (IDT) and their target sequences were summarized in table S4.

For each species, host plants were placed inside the adult cages in the morning for 30-60min to collect eggs. Eggs were soaked in 5% benzalkonium chloride (Sigma-Aldrich) for 20s, rinsed with water, dried and aligned onto 1mm-wide double-sided strips. We used a final concentration of 500ng/µL sgRNA and 500ng/µL Cas9 (UC Berkeley QB3 MacroLab) in the injection mixture and the mixture was incubated at room temperature for 10min before injection. Microinjection was performed using glass capillary needles. Injected eggs were placed back to the growth chamber, and new hatchlings were shifted to artificial diet or host plants to continue development.

### HCR

HCR was performed based on the established protocols^59-61^ with modifications. Probes were designed for *WntA, fz2, pdm3, ivory*, and *Optix* across five nymphalid butterflies. Probes for *WntA* and *pdm3* in *V. cardui, A. incarnata*, and *D. plexippus* were designed in the exons where the corresponding sgRNAs are not located to avoid direct disruption on HCR signals by CRISPR-induced mutagenesis. For *ivory*, probes were designed just downstream of the conserved TSS^39^. All probe DNA oligos were ordered from IDT and their sequences were listed in Data S1.

Pupal wings were dissected at the precisely staged developmental timepoints (table S2) in 1x PBS and fixed in 1x PBST supplemented with 4% formaldehyde for 15-25min, with longer fixation for older wings at room temperature. Wings were washed with PBST and permeabilized with detergent solution^61^ for 10-20min, with longer incubation for older wings. For 8% *A. incarnata* and 6% *D. plexippus* wings, a 3min incubation in detergent was sufficient. Wings were washed first with 1x PBST followed by washes in 5x SSCT. Afterward, wings were transferred to probe hybridization buffer (Molecular Instruments), stored at 4°C temporally or directly proceeded to probe hybridization. Probe hybridization involved incubation in a solution containing 5μL of 100μM probe set mix against each gene in 1mL of probe hybridization buffer for 12-24h at 37°C, followed by washes with probe wash buffer (Molecular Instruments) for 1h at 37°C. Wings were then washed with 5x SSCT and incubated in amplification buffer (Molecular Instruments) for 30min at room temperature. For the chain reaction, wings were incubated in amplification buffer with HCR hairpins (Molecular Instruments) for 4h, followed by washes in 5x SSCT. Finally, wings were mounted on glass slides and cover slips with mounting media. The ventral wing surface was imaged under a Keyence BZ-X810 fluorescence microscope. Images were processed using the BZ-X800 Analyzer software.

We adopted a multiplexing method to remove previous HCR signals and re-stain the same wing tissue multiple rounds with different probe sets whenever necessary^61^. In brief, tissues from the previous staining round were removed from the slides, washed with 5x SSCT at room temperature, then washed with a signal removal buffer 1^61^ at 37°C. HCR signals were then removed by incubating in 400µL of signal removal buffer 1 with 6µL DNaseI at 37°C for one hour. Reaction was terminated by washing in a signal removal buffer 2^61^ at 37°C, followed by washes in signal removal buffer 1 at 37°C. Tissues were then moved to room temperature, washed with 5x SSCT, and transferred to probe hybridization buffer to proceed the next round of staining. For HCR staining, number of biological replicates for each gene, under each genetic background, in each species, was summarized in table S3.

### DNA/RNA extraction

WT and mKO pupal hindwings were HCR stained, imaged, and washed in PBST, flash frozen and stored at -80°C until all the samples were collected. For WT, *WntA* mKO, and *pdm3* mKO pupal wings in *V. cardui* (8% and 17%), *A. incarnata* (13%), and *D. plexippus* (13%), genomic DNA and total RNA were co-extracted using Quick-DNA/RNA Microprep Plus Kit with a pre-digestion and de-crosslinking step. Wings were first digested in 200µL in-house tissue digestion buffer (0.5% (w/v) SDS in 1x TE, pH 8.0, with 10µL proteinase K) at 55°C for 1h and subsequently incubated at 65°C for 15min for de-crosslinking. Equal amount of DNA/RNA Lysis Buffer was added to the reaction, and the rest of the DNA/RNA purification was performed following the manufacturer’s instructions. DNase treatment was included in RNA purification. For *ivory* mKO pupal wings in *V. cardui* (8% and 17%), *pdm3* mKO pupal wings in *A. incarnata* (25%), and *pdm3* mKO adult wings in *A. incarnata* and *D. plexippus*, genomic DNA was extracted using E.Z.N.A Tissue DNA Kit. The resulting genomic DNA from HCR-stained and adult wings was used for genotyping via Sanger sequencing, and total RNA from HCR-stained wings was used for qPCR.

### Genotyping

Genotyping primers were designed to amplify the genomic region spanning each sgRNA cut site (table S4). PCR products were purified using ThermoFisher GeneJET PCR Purification Kit and sent for Sanger sequencing by Azenta. CRISPR guide-induced indel rates were analyzed using the EditCo ICE Analysis tool v4.2.

### qPCR

HCR-stained WT wings and mKO wings with high indel rates (most over 80-90%) were used for qPCR (table S6). 1µg total RNA was reverse-transcribed with oligo-dT using RevertAid First Strand cDNA Synthesis Kit. For *WntA* and *pdm3*, qPCR primers were designed in the exons where the sgRNAs are not located to avoid direct disruption on amplification signals by CRISPR-induced mutagenesis. qPCR was performed using KAPA SYBR FAST Universal Kit. The qPCR thermal cycling program was set as 40 cycles of 95°C for 10s and 60°C for 30s. *Elongation factor 1-alpha* (*EF-1α*) was used as an internal control. For each condition, 3-6 biological replicates were included, and for each biological replicate, three technical replicates were included. Expression data was analyzed using the 2^-ΔΔCt^ method. Melting curves were generated after each qPCR run and checked to ensure unique amplification. All qPCR primers were listed in table S7.

### Statistical analysis

Two-tailed Student’s t-test was used to compare the gene expression levels across WT and mKO wings at each developmental stage in each species. We used p<0.05 as the threshold to infer significance.

