## Supplementary Materials for "Evolutionary rewiring of regulatory topology refines butterfly wing patterns"

### Supplementary Texts

#### Supplementary text 1

To examine whether scale-color master regulators could feedback on the patterning process, we also disrupted the conserved *ivory* transcription start site (TSS) across *V. cardui*, *A. incarnata*, and *D. plexippus*, and checked for corresponding effects on *WntA* and *pdm3* expression. In *V. cardui*, as visible effects on *ivory* expression, a marker to infer *ivory* mKO effect, can only be detected post 17% when *WntA* and *pdm3* are barely expressed, we stained *WntA* and *pdm3* in *ivory* mKO wings at both 8% and 17%, and subsequently genotyped the wings. Confirmed *ivory* mKO mutants showed no visible changes in the expression of *WntA* or *pdm3* at either stage (fig. S9. table S7). In *A. incarnata* and *D. plexippus*, we observed significant reduction in *ivory* expression across *ivory* mKO mutants at 13% (fig. S16-17). No visible changes in the expression of *WntA* or *pdm3* were observed in near-complete *ivory* mKO wings or across *ivory* WT-KO boundaries in the two species (fig. S16-17). This suggests that master color regulators, at least for *ivory*, exhibit minimal feedback on the upstream patterning machineries.

#### Supplementary text 2.

Aligned with adult phenotypes, when *WntA* was disrupted in *D. plexippus*, *ivory* expression was depleted across the wing but remained unchanged around the veins. Meanwhile, *Optix* expression domain remained largely unchanged with slightly less sharp boundaries, and its expression level was visibly unchanged across *WntA* WT-KO boundaries in mKOs (fig. 3D, S13). Of note, *ivory* regulates both orange and black coloration[38], while the role of *Optix* in coloration has not yet been investigated in this species. This suggests that WntA signaling primarily promotes scale coloration in the intervein orange regions and the marginal black bands by activating *ivory*, while it only finetunes the spatial domains of *Optix* in the orange intervein regions and marginal spots. Complementary signaling pathways other than WntA might play a more predominant role in defining the intervein orange patterns and the marginal black bands with spots in this species (fig. 3D, S13).

### Supplementary Figures

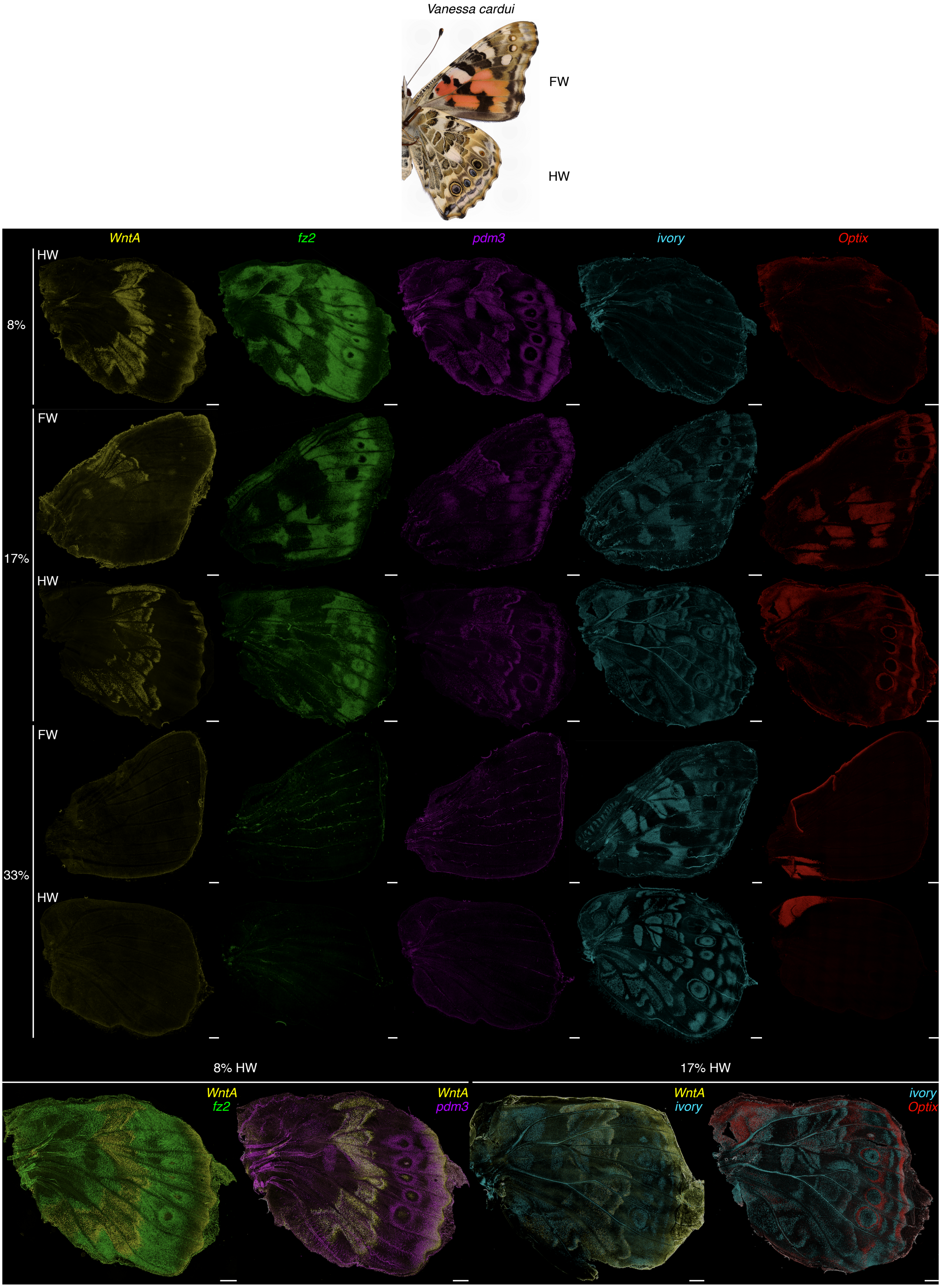


**Fig. S1. Developmental expression dynamics of key butterfly color pattern genes in *V. cardui*.** Adult ventral surfaces and HCR staining on the ventral surfaces of pupal wings are shown. Percentages refer to pupal developmental stages. Numbers of replicates are summarized in table S3. Scale bar, 500 microns.

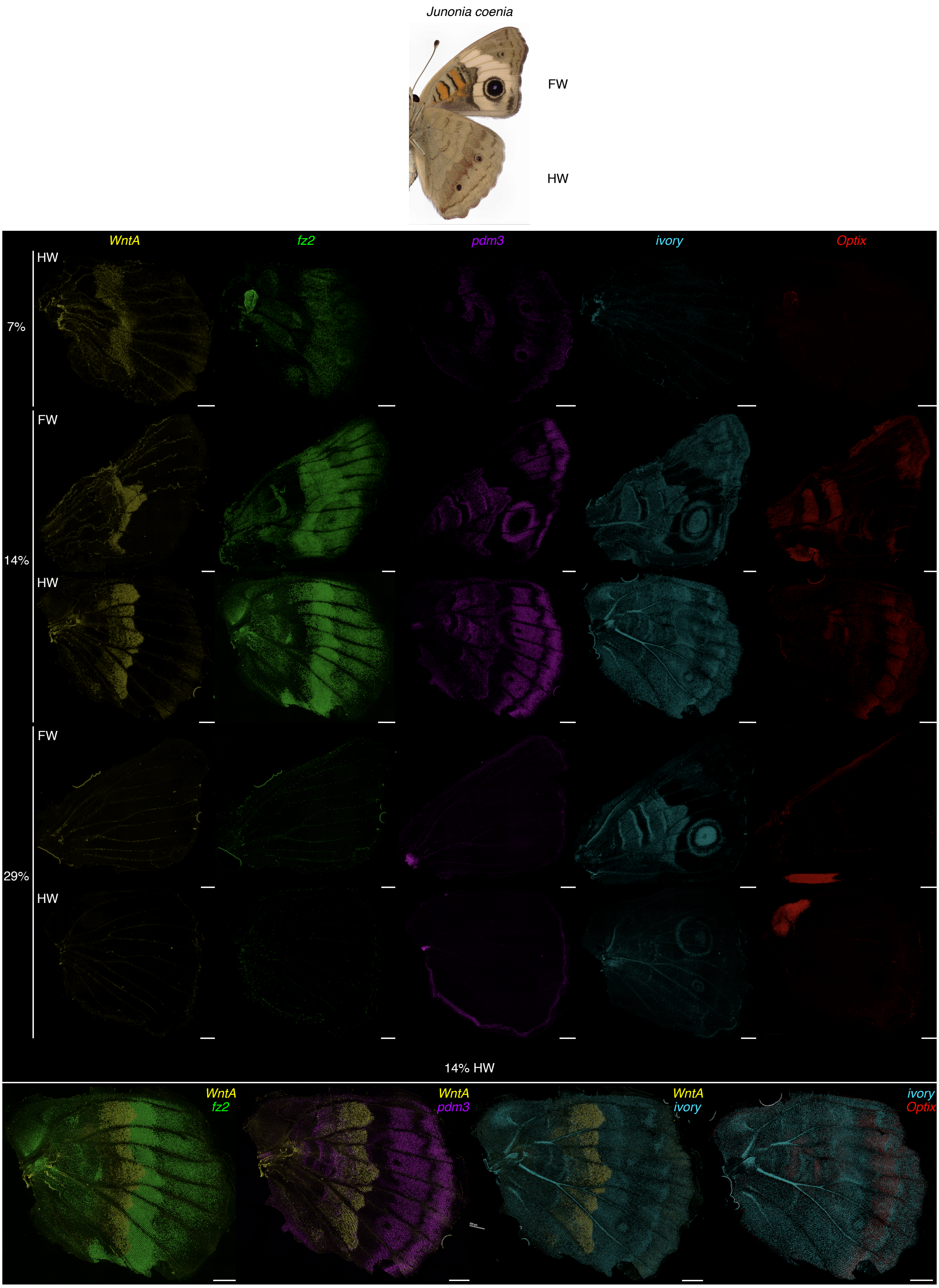


**Fig. S2. Developmental expression dynamics of key butterfly color pattern genes in *J. coenia*.** Adult ventral surfaces and HCR staining on the ventral surfaces of pupal wings are shown. Percentages refer to pupal developmental stages. Numbers of replicates are summarized in table S3. Scale bar, 500 microns.

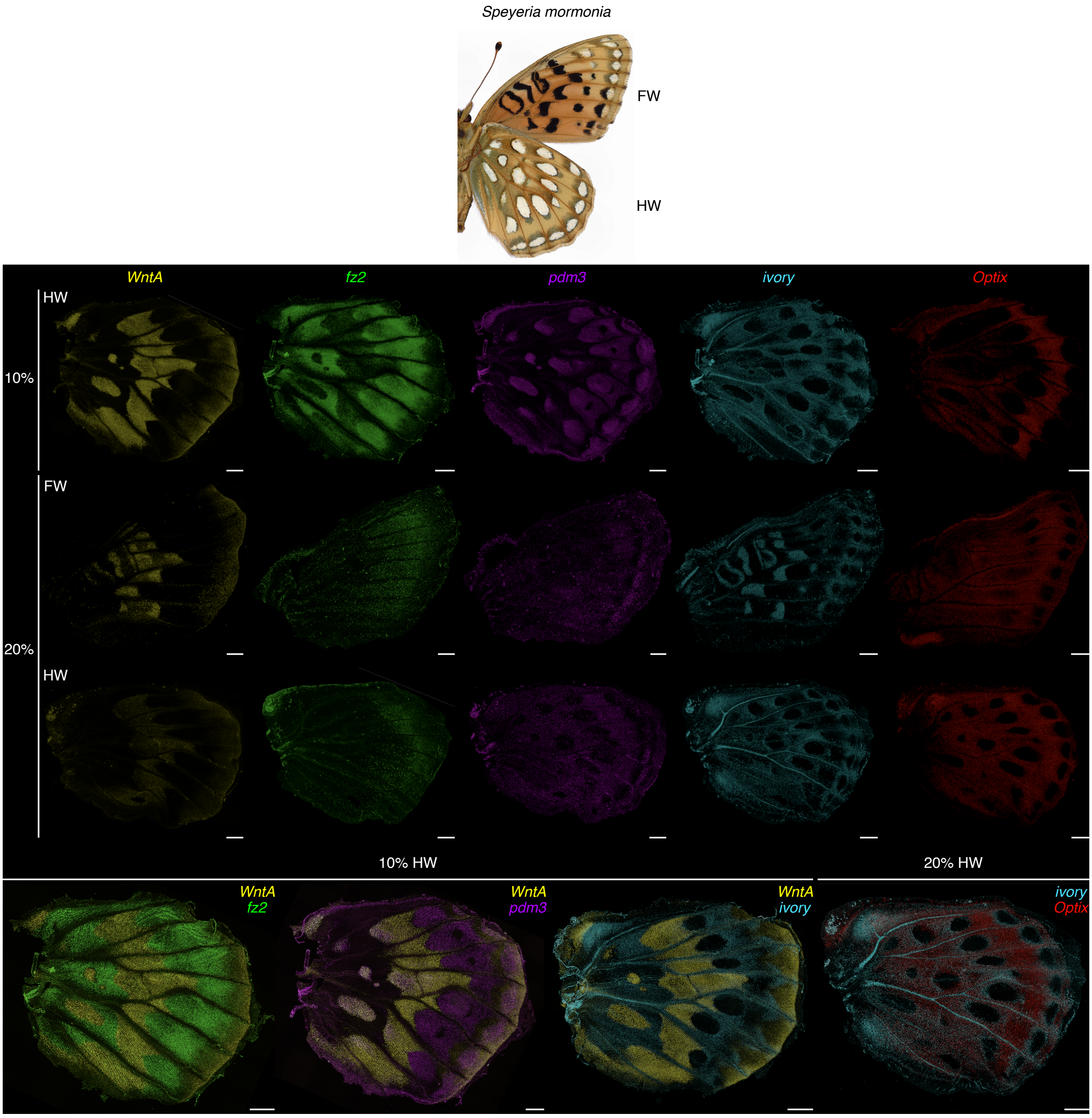


**Fig. S3. Developmental expression dynamics of key butterfly color pattern genes in *S. mormonia*.** Adult ventral surfaces and HCR staining on the ventral surfaces of pupal wings are shown. Percentages refer to pupal developmental stages. Numbers of replicates are summarized in table S3. Scale bar, 500 microns.

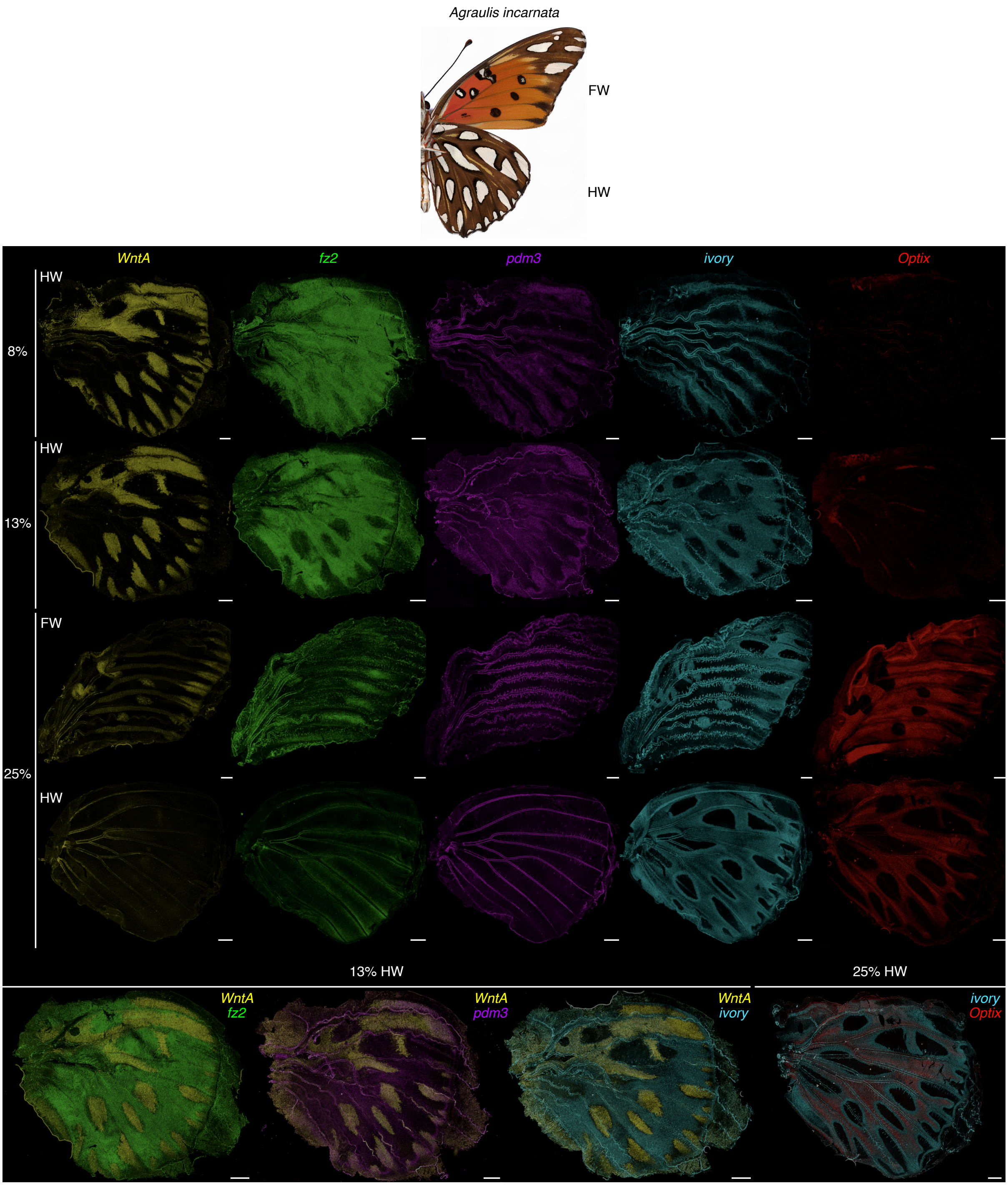


**Fig. S4. Developmental expression dynamics of key butterfly color pattern genes in *A. incarnata*.** Adult ventral surfaces and HCR staining on the ventral surfaces of pupal wings are shown. Percentages refer to pupal developmental stages. Numbers of replicates are summarized in table S3. Scale bar, 500 microns.

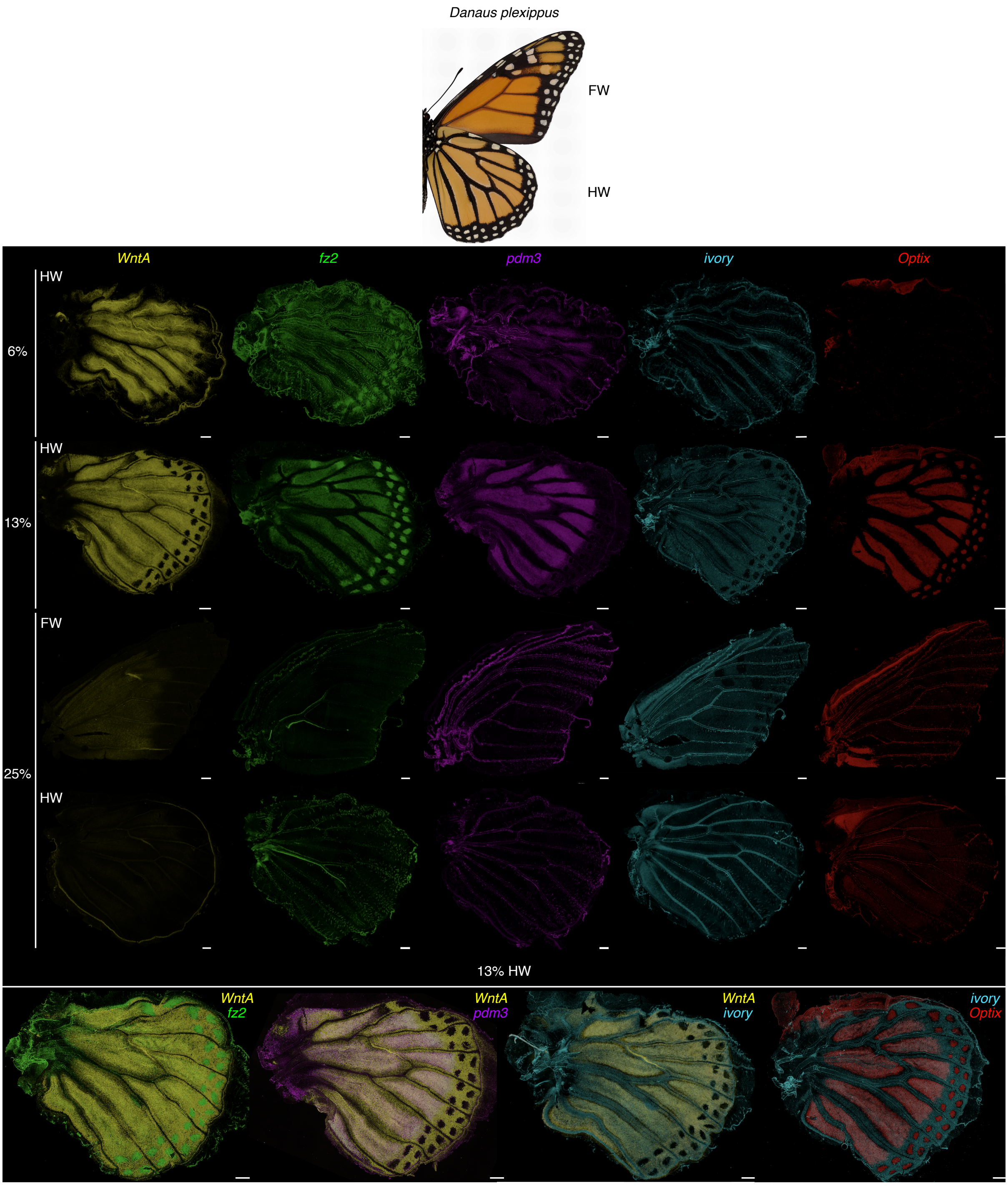


**Fig. S5. Developmental expression dynamics of key butterfly color pattern genes in *D. plexippus*.** Adult ventral surfaces and HCR staining on the ventral surfaces of pupal wings are shown. Percentages refer to pupal developmental stages. Numbers of replicates are summarized in table S3. Scale bar, 500 microns.

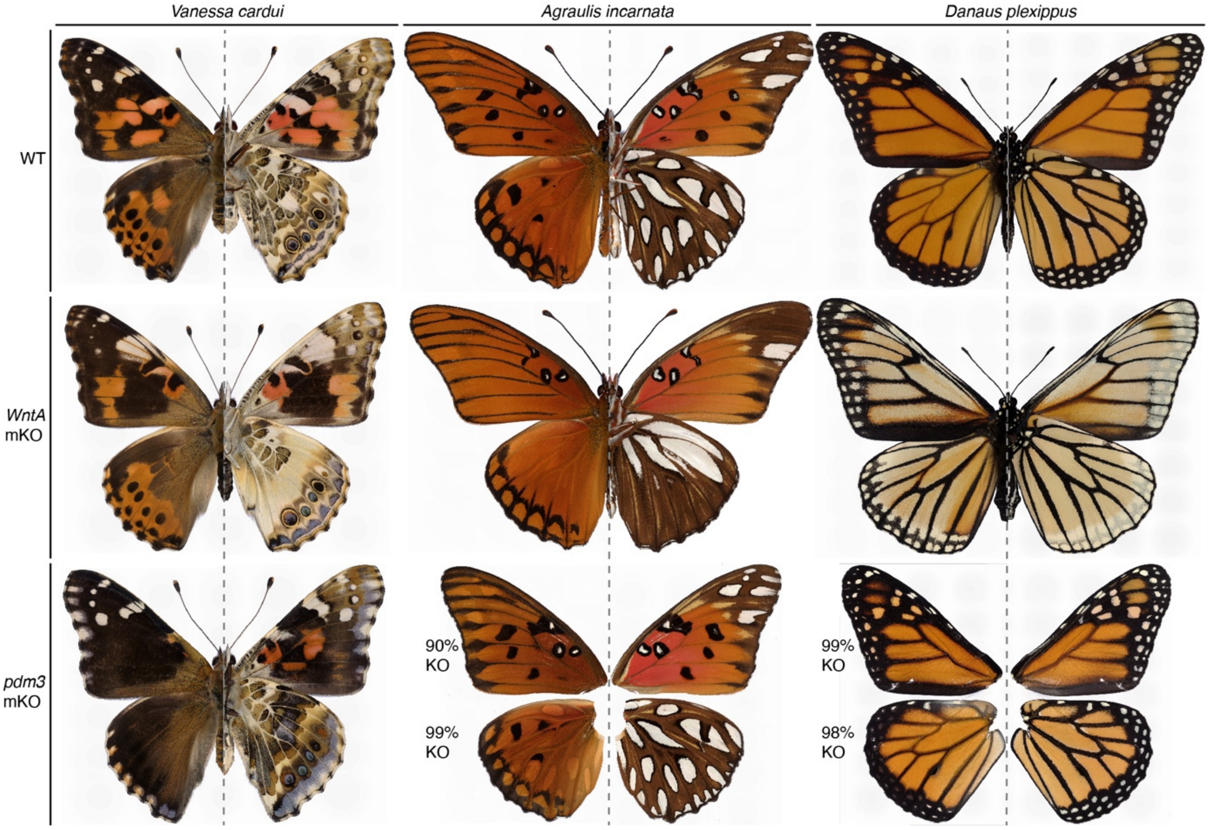


**Fig. S6. WT, *WntA* mKO, and *pdm3* mKO adult wing phenotypes across three representative nymphalids.** Dorsal (left) and ventral (right) surfaces, separated by dotted lines, of WT and near-complete mKO mutants of each species are shown. For mutant wings without visible color pattern changes, wings were genotyped via Sanger sequencing and knockout (KO) rates were shown as indel rates.

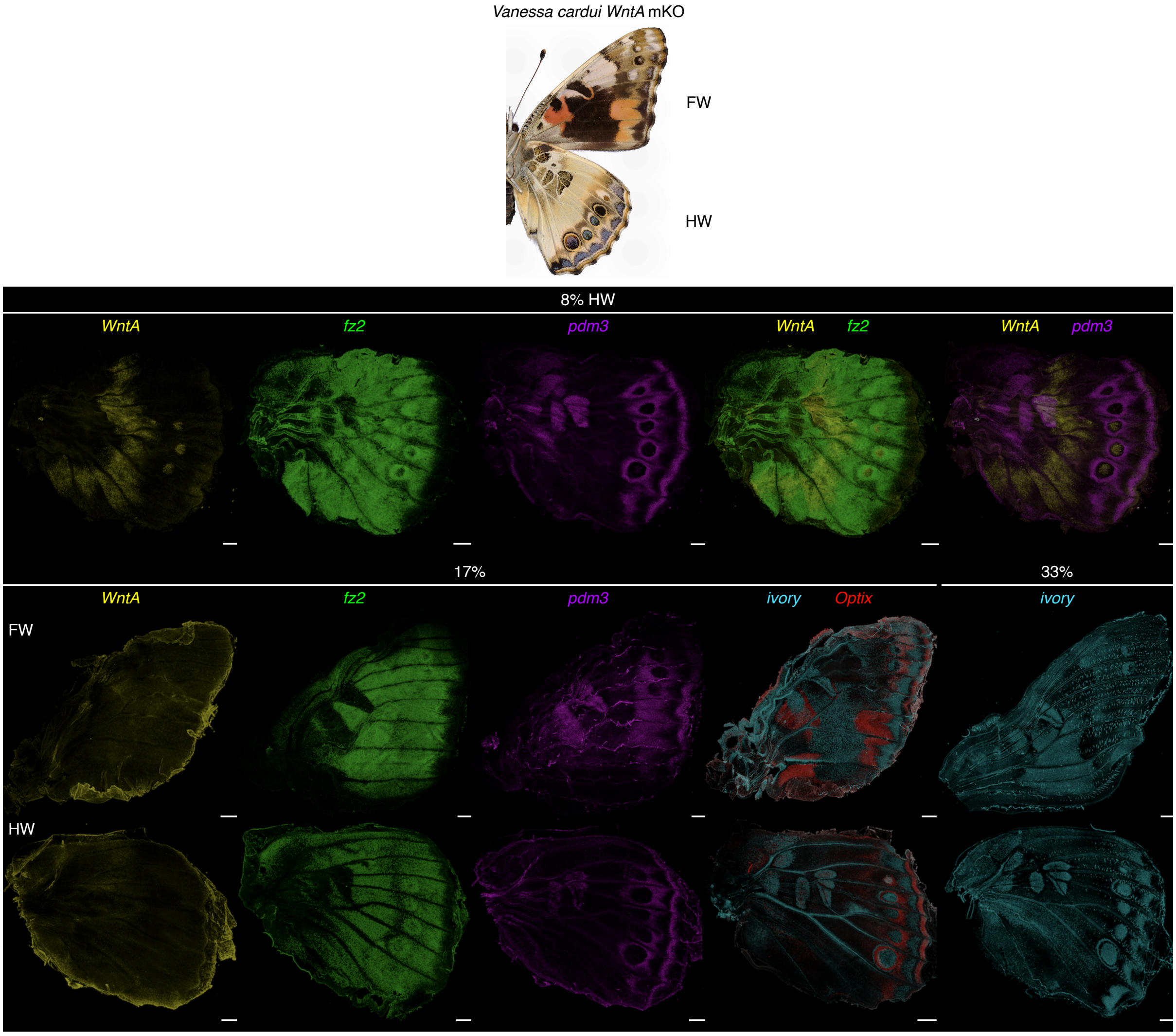


**Fig. S7. Expression of color pattern genes in *WntA* mKO mutants in *V. cardui*.** Near-complete mKO adult ventral surfaces and HCR staining on the ventral surfaces of near-complete mKO pupal wings are shown. Percentages refer to pupal developmental stages. Numbers of replicates are summarized in table S3. Scale bar, 500 microns.

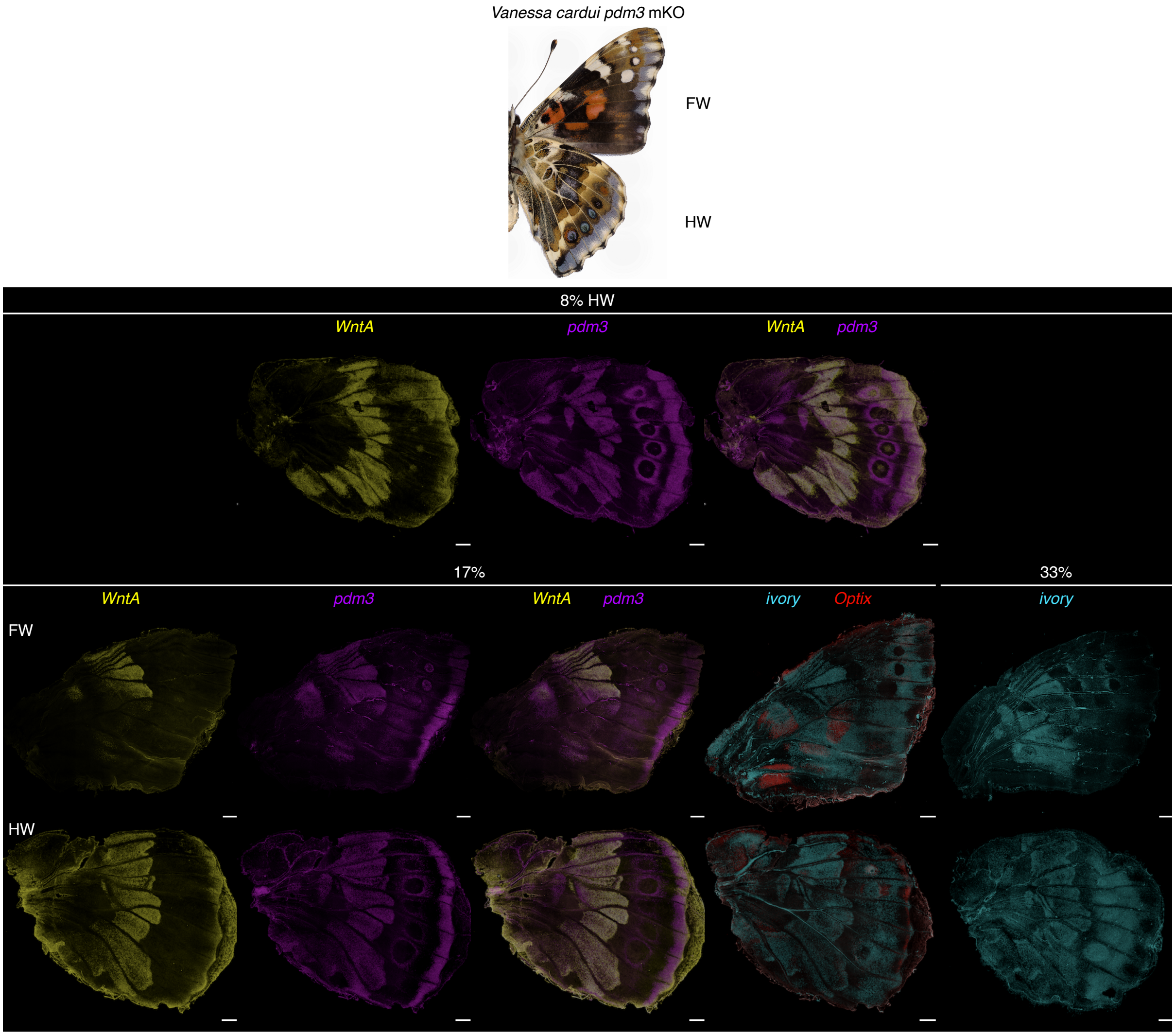


**Fig. S8. Expression of color pattern genes in *pdm3* mKO mutants in *V. cardui*.** Near-complete mKO adult ventral surfaces and HCR staining on the ventral surfaces of near-complete mKO pupal wings are shown. Percentages refer to pupal developmental stages. Numbers of replicates are summarized in table S3. Scale bar, 500 microns.

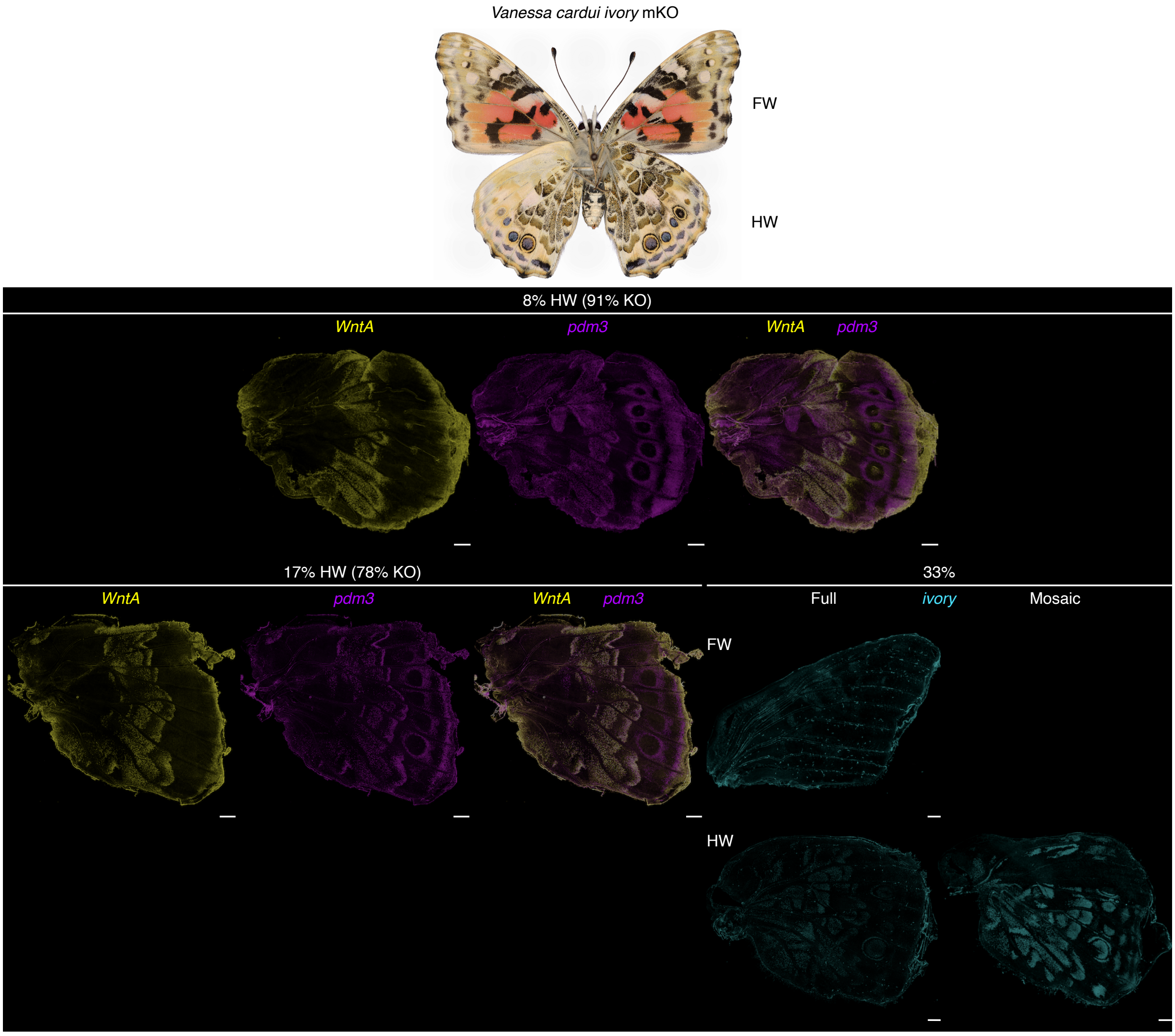


**Fig. S9. Expression of color pattern genes in *ivory* mKO mutants in *V. cardui*.** *ivory* mKO adult ventral surfaces exhibit different shades of reduced melanization. For pupal wings at early stages when *ivory* expression is too low to infer its mKO effects, wings were genotyped via Sanger sequencing, and knockout (KO) rates were shown as indel rates. Percentages refer to pupal developmental stages. Numbers of replicates are summarized in table S3. Scale bar, 500 microns.

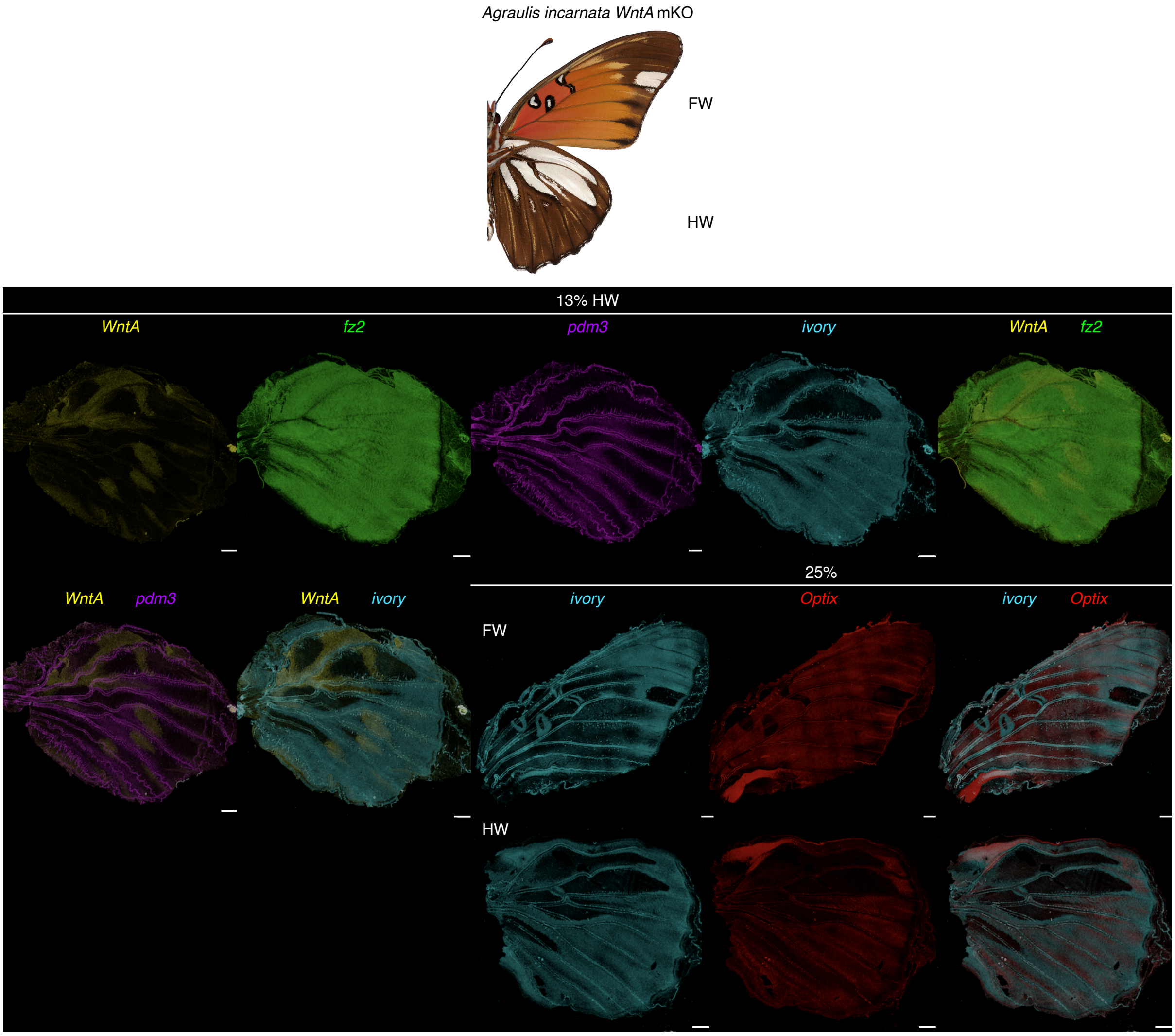


**Fig. S10. Expression of color pattern genes in *WntA* mKO mutants in *A. incarnata*.** Near-complete mKO adult ventral surfaces and HCR staining on the ventral surfaces of near-complete mKO pupal wings are shown. Percentages refer to pupal developmental stages. Numbers of replicates are summarized in table S3. Scale bar, 500 microns.

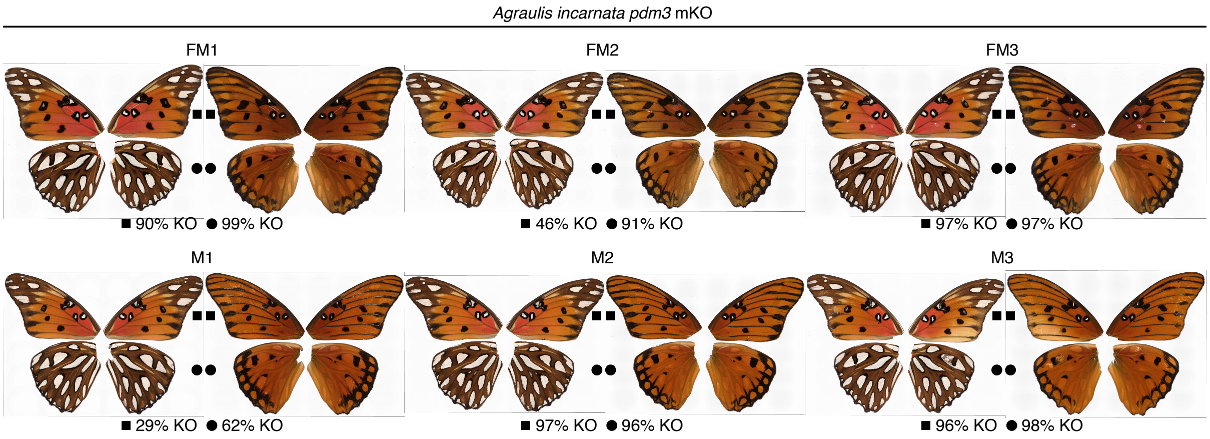


**Fig. S11. *pdm3* mKO mutants show no visibly detectable color pattern changes in *A. incarnata*.** Ventral (left) and dorsal (right) surfaces of adult *pdm3* mKO mutants were shown. Wings without visible color pattern changes, denoted by squares (forewing) and circles (hindwing), were genotyped via Sanger sequencing and knockout (KO) rates were shown as indel rates.

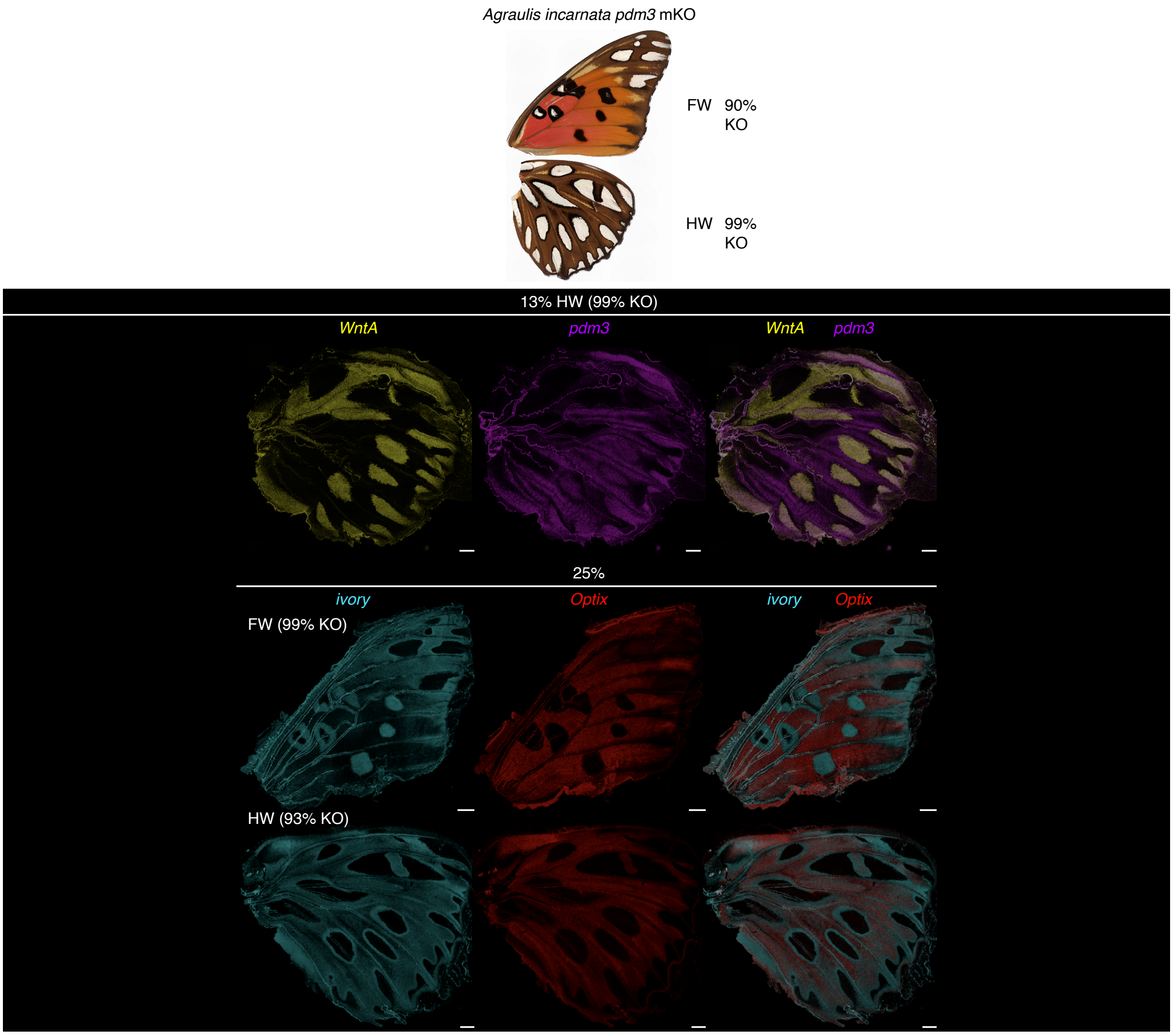


**Fig. S12. Expression of color pattern genes in *pdm3* mKO mutants in *A. incarnata*.** Adult/pupal wings without visible color/expression changes were genotyped via Sanger sequencing and knockout (KO) rates were shown as indel rates. Percentages refer to pupal developmental stages. Numbers of replicates are summarized in table S3. Scale bar, 500 microns.

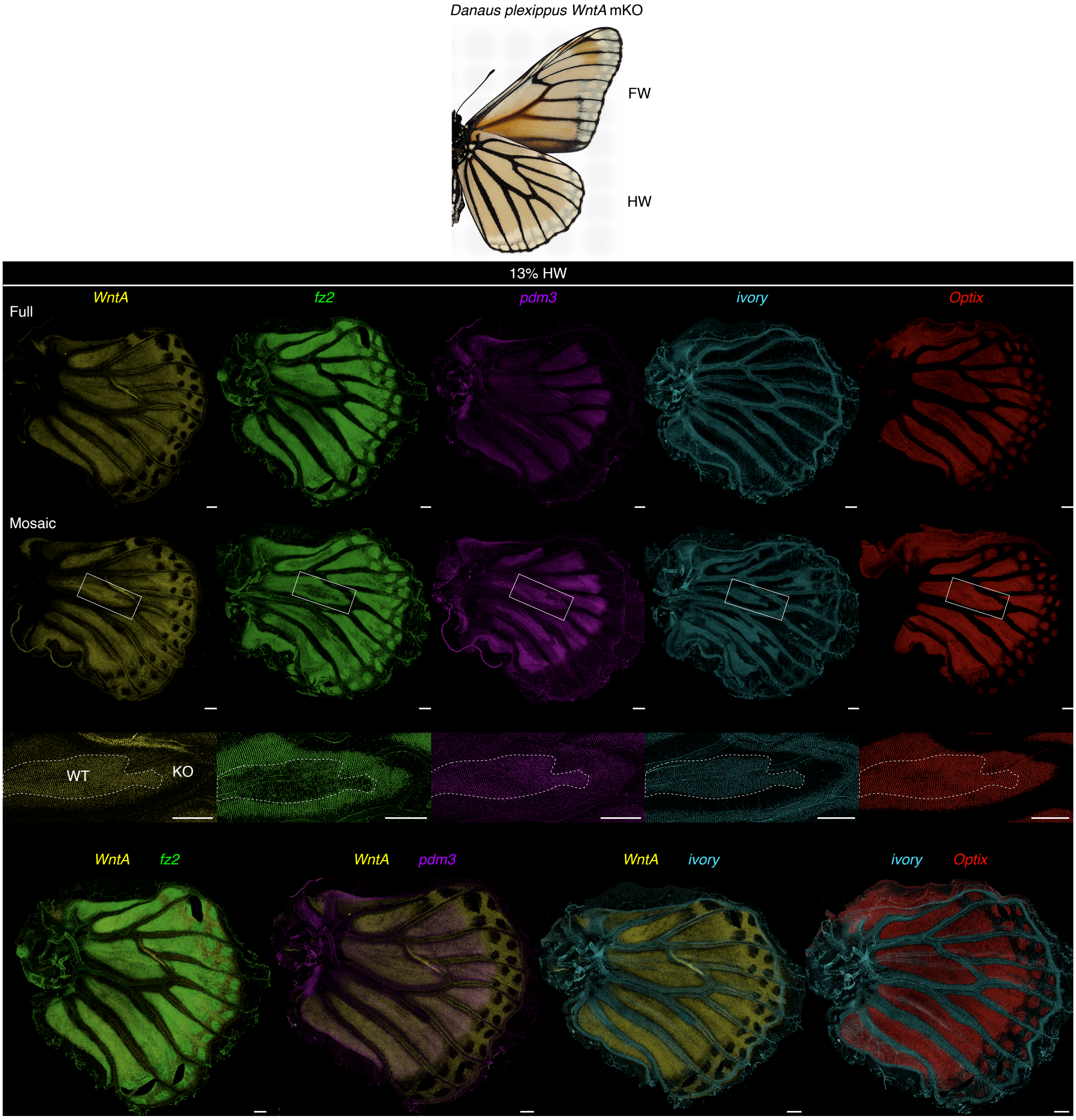


**Fig. S13. Expression of color pattern genes in *WntA* mKO mutants in *D. plexippus*.** Near-complete mKO adult ventral surfaces and HCR staining on the ventral surfaces of both full and mosaic mutant pupal wings are shown. Non-binary changes in expression levels, if any, can be detected across WT-KO boundaries (dotted curves) in the mosaic mutant wings. Percentages refer to pupal developmental stages. Numbers of replicates are summarized in table S3. Scale bar, 500 microns.

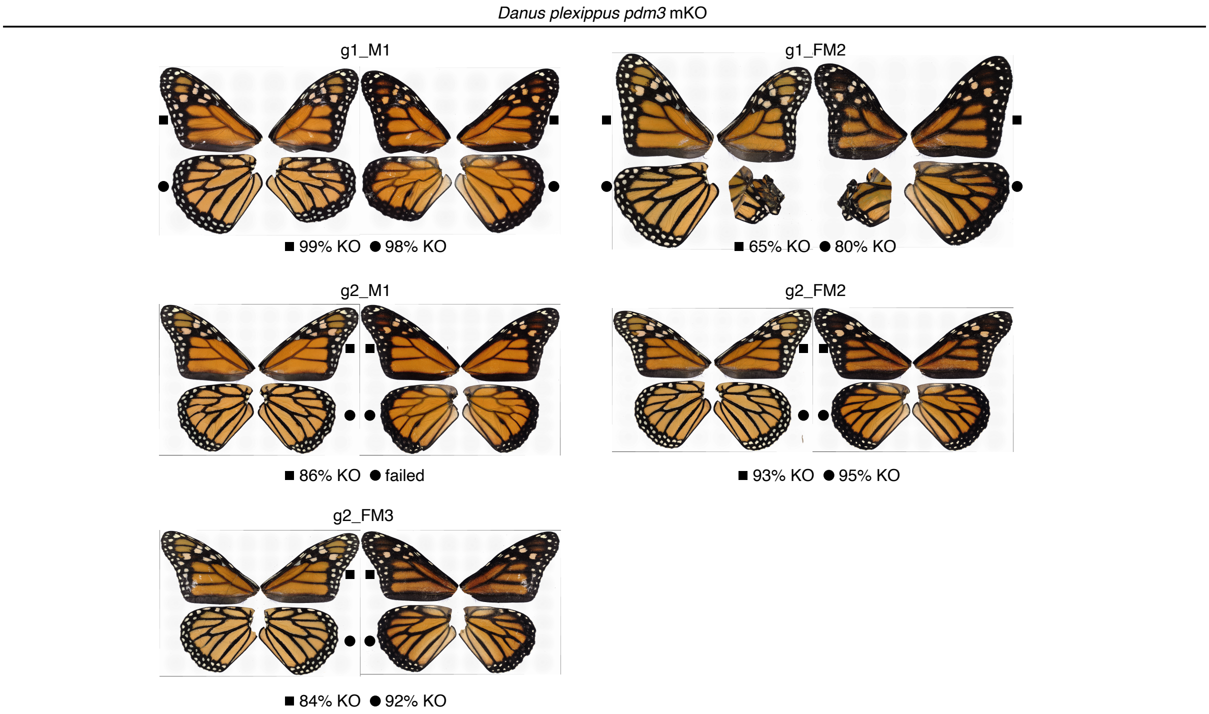


**Fig. S14. *pdm3* mKO mutants show no visibly detectable color pattern changes in *D. plexippus*.** Ventral (left) and dorsal (right) surfaces of adult *pdm3* mKO mutants were shown. Wings without visible color pattern changes, denoted by squares (forewing) and circles (hindwing), were genotyped via Sanger sequencing and knockout (KO) rates were shown as indel rates. This was validated across two individual guide RNAs, g1 and g2.

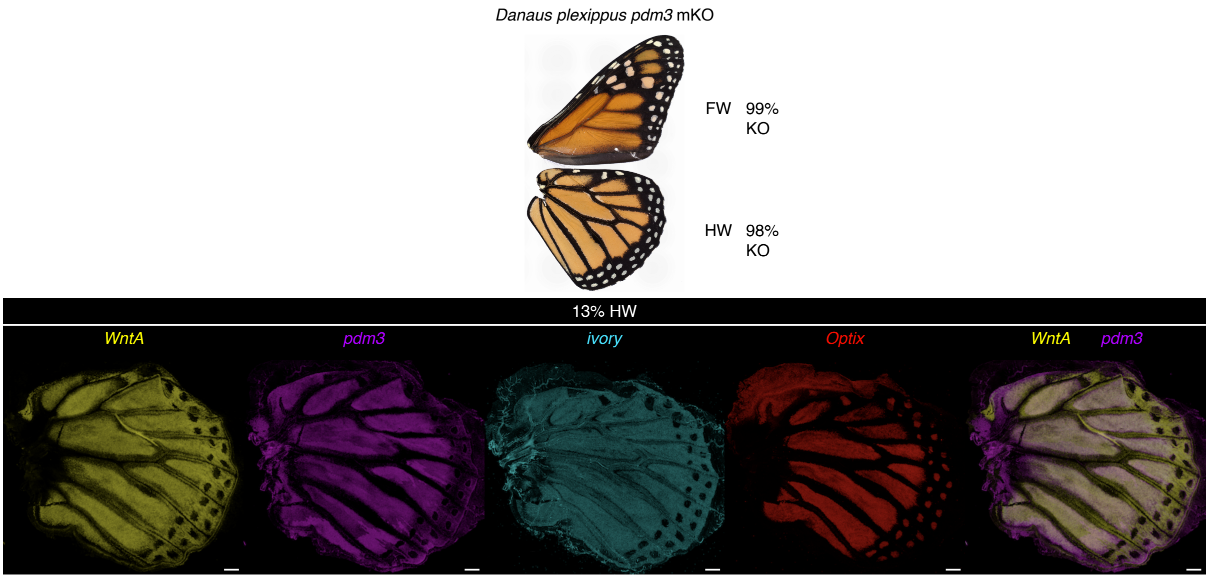


**Fig. S15. Expression of color pattern genes in *pdm3* mKO mutants in *D. plexippus*.** Adult wings without visible color pattern changes were genotyped via Sanger sequencing and knockout (KO) rates were shown as indel rates. Boundaries with different *pdm3* expression levels were observed across the mKO pupal wings, without visible changes in the expression of other color pattern genes across these boundaries. Percentages refer to pupal developmental stages. Numbers of replicates are summarized in table S3. Scale bar, 500 microns.

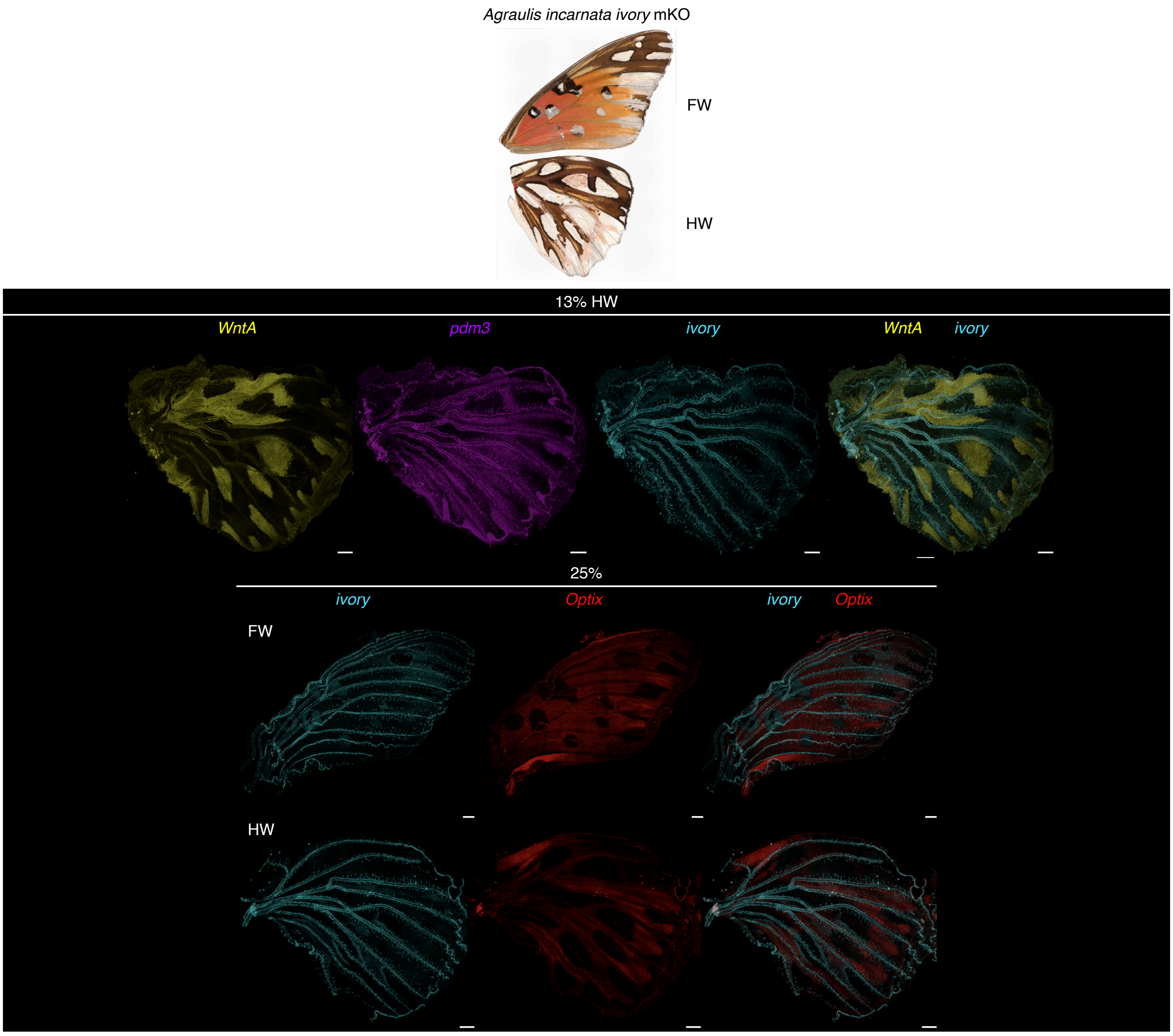


**Fig. S16. Expression of color pattern genes in *ivory* mKO mutants in *A. incarnata*.** *ivory* mKO adult ventral surfaces exhibit different shades of reduced melanization and mKO pupal wings exhibit reduced or almost undetectable levels of *ivory* expression, with no visible changes in the expression of other color pattern genes. Percentages refer to pupal developmental stages. Numbers of replicates are summarized in table S3. Scale bar, 500 microns.

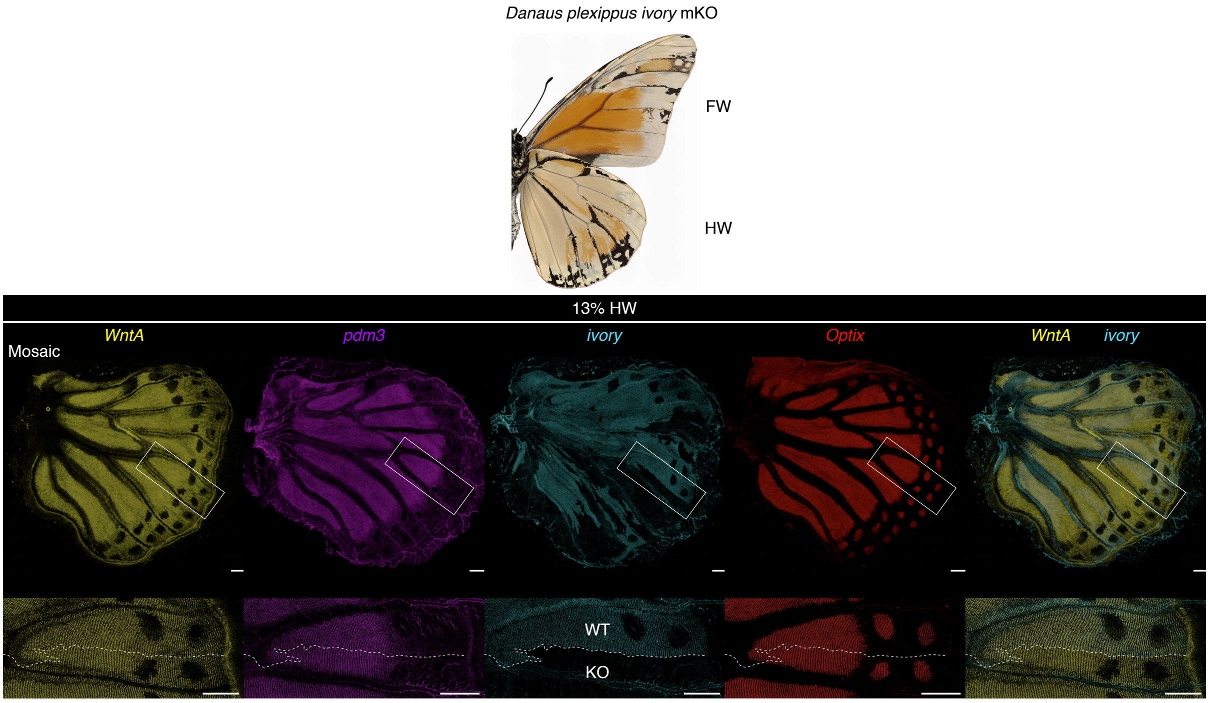


**Fig. S17. Expression of color pattern genes in *ivory* mKO mutants in *D. plexippus*.** *ivory* mKO adult ventral surfaces exhibit different shades of reduced melanization and mKO pupal wings exhibit WT-KO boundaries with presence and absence of *ivory* expression (dotted curves), without visible changes in the expression of other color pattern genes across these boundaries. Percentages refer to pupal developmental stages. Numbers of replicates are summarized in table S3. Scale bar, 500 microns.

### Supplementary Tables

**Table S1. Animal husbandry.**

| Species | Source | Host plant/Artificial diet |
|---|---|---|
| *V. cardui* | #144070, Carolina Biological Supply | Multiple species diet, Southland Products |
| *J. coenia* | Robert Reed lab, Cornell University | Artificial diet described in H. F. Nijhout *Dev. Biol.* (1980) |
| *S. mormonia* | Carol Boggs lab, University of South Carolina | *Viola sororia* |
| *A. incarnata* | Kitty Bryan, FL, USA<br>SK Butterflies, FL, USA | *Passiflora incarnata* |
| *D. plexippus* | Monarch Watch, KS, USA<br>Rose Franklin's Perennial & Butterfly Farm, PA, USA | *Asclepias curassavica* |

**Table S2. Pupal development time.**

| Species | % pupal development (hours post-pupation) | Total (days)* |
|---|---|---|
| *V. cardui* | 8% (12h), 17% (24h), 33% (48h) | 6.0 |
| *J. coenia* | 7% (12h), 14% (24h), 29% (48h) | 6.9 |
| *S. mormonia* | 10% (24h), 20% (48h) | 9.9 |
| *A. incarnata* | 8% (16h), 13% (24h), 25% (48h) | 8.0 |
| *D. plexippus* | 6% (12h), 13% (24h), 25% (48h) | 7.9 |

*: Under 27°C, 16:8 day:night cycle, 60% relative humidity

**Table S3. Number of biological replicates for HCR staining.**

| Species | Genotype | Pupal development % | Wing** | Number of wings | Number of replicates | | | | |
|---|---|---|---|---|---|---|---|---|---|
| | | | | | *WntA* | *fz2* | *pdm3* | *ivory* | *Optix* |
| *V. cardui* | WT | 8% | HW | 4 | 4 | 4 | 4 | 4 | 4 |
| | | 17% | FW | 7 | 7 | 6 | 6 | 7 | 4 |
| | | 17% | HW | 16 | 13 | 4 | 4 | 16 | 6 |
| | | 33% | FW | 3 | 3 | 3 | 3 | 3 | 3 |
| | | 33% | HW | 4 | 4 | 4 | 4 | 4 | 4 |
| | *WntA* mKO | 8% | HW | 10 | 10 | 2 | 10 | | |
| | | 17% | FW | 6 | 6 | 5 | 4 | 6 | 6 |
| | | 17% | HW | 11 | 11 | 7 | 7 | 7 | 7 |
| | | 33% | FW | 4 | | | | 4 | |
| | | 33% | HW | 3 | | | | 3 | |
| | *pdm3* mKO | 8% | HW | 18 | 11 | | 18 | | |
| | | 17% | FW | 4 | 4 | | 4 | 4 | 4 |
| | | 17% | HW | 8 | 8 | | 8 | 2 | 2 |
| | | 33% | FW | 4 | | | | 4 | |
| | | 33% | HW | 3 | | | | 3 | |
| | *ivory* mKO | 8% | HW | 11 | 11 | | 11 | | |
| | | 17% | FW | 0 | | | | | |
| | | 17% | HW | 9 | 9 | | 9 | | |
| | | 33% | FW | 4 | | | | 4 | |
| | | 33% | HW | 3 | | | | 3 | |
| *J.coenia* | WT | 7% | HW | 1 | 1 | 1 | 1 | 1 | 1 |
| | | 14% | FW | 2 | 2 | 2 | 2 | 2 | 2 |
| | | 14% | HW | 3 | 3 | 3 | 3 | 3 | 3 |
| | | 29% | FW | 2 | 2 | 2 | 2 | 2 | 2 |
| | | 29% | HW | 3 | 3 | 3 | 3 | 3 | 3 |
| *S. mormonia* | WT | 10% | HW | 2 | 2 | 2 | 2 | 2 | 2 |
| | | 20% | FW | 2 | 2 | 2 | 2 | 2 | 2 |
| | | 20% | HW | 2 | 2 | 2 | 2 | 2 | 2 |
| *A. incarnata* | WT | 8% | HW | 2 | 2 | 2 | 2 | 2 | 2 |
| | | 13% | HW | 10 | 10 | 6 | 6 | 2 | 2 |
| | | 25% | FW | 3 | 3 | 2 | 2 | 3 | 3 |
| | | 25% | HW | 4 | 4 | 2 | 2 | 4 | 2 |
| | *WntA* mKO | 13% | HW | 12 | 12 | 2 | 8 | 8 | |
| | | 25% | FW | 4 | | | | 4 | 4 |
| | | 25% | HW | 4 | | | | 4 | 4 |
| | *pdm3* mKO | 13% | HW | 14 | 14 | | 14 | | |
| | | 25% | FW | 5 | | | | 5 | 5 |
| | | 25% | HW | 6 | | | | 6 | 6 |
| | *ivory* mKO | 13% | HW | 10 | 10 | | 2 | 10 | |
| | | 25% | FW | 1 | | | | 1 | 1 |
| | | 25% | HW | 3 | | | | 3 | 3 |
| *D. plexippus* | WT | 6% | HW | 4 | 4 | 4 | 2 | 4 | 4 |
| | | 13% | HW | 8 | 6 | 3 | 3 | 4 | 6 |
| | | 25% | FW | 4 | 2 | 2 | 3 | 2 | 2 |
| | | 25% | HW | 4 | 2 | 2 | 3 | 2 | 2 |
| | *WntA* mKO | 13% | HW | 8 | 8 | 3 | 3 | 8 | 3 |
| | *pdm3* mKO* | 13% | HW | 10 | 10 | | 10 | 2 | 2 |
| | *ivory* mKO | 13% | HW | 10 | 10 | | 5 | 10 | 5 |

*: sgRNA used: Dp_pdm3_g1

**: FW, Forewing; HW, hindwing

**Table S4. Guide RNAs and genotyping primers.**

| Species | NCBI Genome | sgRNA | Target sequence | Genotyping primer | Sequence |
|---|---|---|---|---|---|
| *V. cardui* | ASM2240509v1 | Vc_WntA_g1 | GGCAGCATTGGCCCATGCGG | GT_Vc_WntA_F | ACACAACAACAAGCAGCAGA |
| | | | | GT_Vc_WntA_R | CCAATGTCGAGGAGTGCGTT |
| | | | | GT_Vc_WntA_R2 | ACTGCCAATGTCGAGGAGTG |
| | | Vc_pdm3_g1 | CAGCGCTTGAGGCGTGAATG | GT_Vc_pdm3_F | GTGTCCGGGTAATTAATGGCTAAG |
| | | | | GT_Vc_pdm3_F2 | GCGTTCTCAGATTTACTTCTCTAGG |
| | | | | GT_Vc_pdm3_R | ATCCTTCCAGGTACGCGTCG |
| | | | | GT_Vc_pdm3_R2 | ATGCCACTTATCAACCCGGACC |
| | | VcAi_ivory_g1 | CGCGATCTACGCGACGAATG | GT_Vc_ivory_F | TTTGGCAAGGACGTTCGTTG |
| | | | | GT_Vc_ivory_R | GGGACACCATCTCCAATCAACT |
| *A. incarnata* | Avfl.assembly.v1.2 | Ai_WntA_g1 | GGCTCGGGCGTGCAGTCGTG | GT_Ai_WntA_F | GGCTAAATGGGGTGACGTTGTC |
| | | | | GT_Ai_WntA_R | GCACGTAACAATACGTGGTTGC |
| | | Ai_pdm3_g1 | CAGTTGGCTGCCAACCTCCA | GT_Ai_pdm3_F | GTCCACAGTGGCATCTATGT |
| | | | | GT_Ai_pdm3_R | ATGATATTTACCTGGGGCGC |
| | | VcAi_ivory_g1 | CGCGATCTACGCGACGAATG | | |
| *D. plexippus* | MEX_DaPlex | Dp_WntA_g1 | GCTATCCGACAAGATGGTTC | GT_Dp_WntA_F | TGGACCACTTCATTCAGAATGG |
| | | | | GT_Dp_WntA_R | GGTTCAAACCTCGTACATTGCT |
| | | Dp_pdm3_g1 | ACGGACGTCCTTCACGCCGC | GT_Dp_pdm3_F | AAGGTCTATTTTTCACAGGTACG |
| | | | | GT_Dp_pdm3_R | GAGACTATAACCGAATAGACAAG |
| | | Dp_pdm3_g2 | GCGTGCGCCCGCGCACCCTC | GT_Dp_pdm3_F2 | TCCACAAGATCCTAGGCGAG |
| | | | | GT_Dp_pdm3_R2 | GGGTGGGAGTTTAAATAAAGACG |
| | | Dp_ivory_g1 | CGCGATCTCCGCGACGAATG | | |

**Table S5. Genotyping results of HCR-stained wings used for qPCR.**

| Species | sgRNA | Sample ID | Pupal development % | GT primers | Primer used for Sanger sequencing | Indel% |
|---|---|---|---|---|---|---|
| *V. cardui* | Vc_WntA_g1 | WntA_0.5_6 | 8% | GT_Vc_WntA_F and R | GT_Vc_WntA_F | 100% |
| | | WntA_0.5_10 | 8% | GT_Vc_WntA_F and R | GT_Vc_WntA_F | 76% |
| | | WntA_0.5_9 | 8% | GT_Vc_WntA_F and R2 | GT_Vc_WntA_F | 89% |
| | | WntA_1_11 | 17% | GT_Vc_WntA_F and R | GT_Vc_WntA_F | 96% |
| | | WntA_1_4 | 17% | GT_Vc_WntA_F and R2 | GT_Vc_WntA_F | 82% |
| | | WntA_1_9 | 17% | GT_Vc_WntA_F and R2 | GT_Vc_WntA_F | 79% |
| | Vc_pdm3_g1 | pdm_0.5_6 | 8% | GT_Vc_pdm3_F2 and R2 | GT_Vc_pdm3_R2 | 65% |
| | | pdm_0.5_11 | 8% | GT_Vc_pdm3_F and R | GT_Vc_pdm3_F | 70% |
| | | pdm_0.5_13 | 8% | GT_Vc_pdm3_F and R | GT_Vc_pdm3_F | 78% |
| | | pdm_0.5_14 | 8% | GT_Vc_pdm3_F2 and R2 | GT_Vc_pdm3_R2 | 60% |
| | | pdm_1_3 | 17% | GT_Vc_pdm3_F and R | GT_Vc_pdm3_F | 60% |
| | | pdm_1_5 | 17% | GT_Vc_pdm3_F and R | GT_Vc_pdm3_F | 78% |
| | | pdm_1_6 | 17% | GT_Vc_pdm3_F and R | GT_Vc_pdm3_F | 75% |
| | | pdm_1_7 | 17% | GT_Vc_pdm3_F2 and R2 | GT_Vc_pdm3_R2 | 77% |
| *A. incarnata* | Ai_WntA_g1 | WntA_5 | 13% | GT_Ai_WntA_F and R | GT_Ai_WntA_F | 94% |
| | | WntA_7 | 13% | GT_Ai_WntA_F and R | GT_Ai_WntA_F | 94% |
| | | WntA_11 | 13% | GT_Ai_WntA_F and R | GT_Ai_WntA_F | 95% |
| | | WntA_13 | 13% | GT_Ai_WntA_F and R | GT_Ai_WntA_F | 69% |
| | | WntA_14 | 13% | GT_Ai_WntA_F and R | GT_Ai_WntA_F | 72% |
| | Ai_pdm3_g1 | pdm_HW2 | 13% | GT_Ai_pdm3_F and R | GT_Ai_pdm3_F | 99% |
| | | pdm_HW6 | 13% | GT_Ai_pdm3_F and R | GT_Ai_pdm3_F | 98% |
| | | pdm_HW8 | 13% | GT_Ai_pdm3_F and R | GT_Ai_pdm3_F | 97% |
| | | pdm_HW12 | 13% | GT_Ai_pdm3_F and R | GT_Ai_pdm3_F | 91% |
| | | pdm_HW14 | 13% | GT_Ai_pdm3_F and R | GT_Ai_pdm3_F | 99% |
| *D. plexippus* | Dp_WntA_g1 | WntA_HW9 | 13% | GT_Dp_WntA_F and R | GT_Dp_WntA_R | 80% |
| | | WntA_HW10 | 13% | GT_Dp_WntA_F and R | GT_Dp_WntA_R | 73% |
| | | WntA_HW11 | 13% | GT_Dp_WntA_F and R | GT_Dp_WntA_R | 80% |
| | Dp_pdm3_g1 | pdm_g1_HW1 | 13% | GT_Dp_pdm3_F and R | GT_Dp_pdm3_R | 97% |
| | | pdm_g1_HW2 | 13% | GT_Dp_pdm3_F and R | GT_Dp_pdm3_R | 85% |
| | | pdm_g1_HW7L | 13% | GT_Dp_pdm3_F and R | GT_Dp_pdm3_R | 83% |

**Table S6. qPCR primers.**

| Species | Target gene | qPCR primer | Sequence |
|---|---|---|---|
| *V. cardui* | *EF-1a* | qPCR_Vc_EF1a_F | AGCCCCCATACAATGAAGGC |
| | | qPCR_Vc_EF1a_R | AATGGGTACGAAAGCGACGG |
| | *WntA* | qPCR_Vc_WntA_F | TCGTGGTCGAATGCAACGT |
| | | qPCR_Vc_WntA_R | TCACCCACGATCCTTAGTTGTG |
| | *pdm3* | qPCR_Vc_pdm3_F | AATGTCTATCGGGCCTTGGC |
| | | qPCR_Vc_pdm3_R | GCAGGCGAATCACAACCTTC |
| *A. incarnata* | *EF-1a* | qPCR_Ai_EF1a_F | AATGGGTACGAAAGCGACGG |
| | | qPCR_Ai_EF1a_R | GCCTCCATACAGCGAGTCTC |
| | *WntA* | qPCR_Ai_WntA_F | CATGTCGGGCTCCTGTTCTG |
| | | qPCR_Ai_WntA_R | AAGCGCCTTCGTATCTCGTC |
| | *pdm3* | qPCR_Ai_pdm3_F | TGCAAGGCAGAAGGTCATCC |
| | | qPCR_Ai_pdm3_R | CCCTTTCATTTCGTTCCCTAGA |
| *D. plexippus* | *EF-1a* | qPCR_Dp_EF1a_F | GCTCGATTTGCCATCCCTTG |
| | | qPCR_Dp_EF1a_R | GCTGTCGCTTTCGTACCCAT |
| | *WntA* | qPCR_Dp_WntA_F | ACATGACTCGCTCCCTCGTA |
| | | qPCR_Dp_WntA_R | TGGCATGTCTGGTTCCTGTT |
| | *pdm3* | qPCR_Dp_pdm3_F | CAACGTCCCATTCAACAGCG |
| | | qPCR_Dp_pdm3_R | CTTGGTGAGAGCTTGACGGG |

**Table S7. Genotyping results of HCR-stained wings without a marker gene.**

| Species | sgRNA | Sample ID | Pupal development % | GT primers | Primer used for Sanger sequencing | Indel% |
|---|---|---|---|---|---|---|
| *V. cardui* | VcAi_ivory_g1 | ivory_0.5_3 | 8% | GT_Vc_ivory_F and R | GT_Vc_ivory_F | 72% |
| | | ivory_0.5_4 | 8% | GT_Vc_ivory_F and R | GT_Vc_ivory_F | 85% |
| | | ivory_0.5_5 | 8% | GT_Vc_ivory_F and R | GT_Vc_ivory_F | 91% |
| | | ivory_0.5_11 | 8% | GT_Vc_ivory_F and R | GT_Vc_ivory_F | 95% |
| | | ivory_0.5_6 | 8% | GT_Vc_ivory_F and R | GT_Vc_ivory_F | 67% |
| | | ivory_0.5_9 | 8% | GT_Vc_ivory_F and R | GT_Vc_ivory_F | 93% |
| | | ivory_0.5_8 | 8% | GT_Vc_ivory_F and R | GT_Vc_ivory_F | 92% |
| | | ivory_0.5_7 | 8% | GT_Vc_ivory_F and R | GT_Vc_ivory_F | 92% |
| | | ivory_0.5_10 | 8% | GT_Vc_ivory_F and R | GT_Vc_ivory_F | 90% |
| | | ivory_1_3 | 17% | GT_Vc_ivory_F and R | GT_Vc_ivory_F | 73% |
| | | ivory_1_6 | 17% | GT_Vc_ivory_F and R | GT_Vc_ivory_F | 78% |
| | | ivory_1_5 | 17% | GT_Vc_ivory_F and R | GT_Vc_ivory_F | 72% |
| | | ivory_1_4 | 17% | GT_Vc_ivory_F and R | GT_Vc_ivory_F | fail |
| | | ivory_1_7 | 17% | GT_Vc_ivory_F and R | GT_Vc_ivory_R | 60% |
| | | ivory_1_8 | 17% | GT_Vc_ivory_F and R | GT_Vc_ivory_R | fail |
| | | ivory_1_9 | 17% | GT_Vc_ivory_F and R | GT_Vc_ivory_R | 84% |
| *A. incarnata* | Ai_pdm3_g1 | pdm_day2_FW1 | 25% | GT_Ai_pdm3_F and R | GT_Ai_pdm3_F | 94% |
| | | pdm_day2_FW2 | 25% | GT_Ai_pdm3_F and R | GT_Ai_pdm3_F | fail |
| | | pdm_day2_FW3 | 25% | GT_Ai_pdm3_F and R | GT_Ai_pdm3_F | 97% |
| | | pdm_day2_FW4 | 25% | GT_Ai_pdm3_F and R | GT_Ai_pdm3_F | 99% |
| | | pdm_day2_FW5 | 25% | GT_Ai_pdm3_F and R | GT_Ai_pdm3_F | fail |
| | | pdm_day2_HW1 | 25% | GT_Ai_pdm3_F and R | GT_Ai_pdm3_F | 93% |
| | | pdm_day2_HW2 | 25% | GT_Ai_pdm3_F and R | GT_Ai_pdm3_F | 93% |
| | | pdm_day2_HW3 | 25% | GT_Ai_pdm3_F and R | GT_Ai_pdm3_F | 94% |
| | | pdm_day2_HW4 | 25% | GT_Ai_pdm3_F and R | GT_Ai_pdm3_F | 96% |
| | | pdm_day2_HW5 | 25% | GT_Ai_pdm3_F and R | GT_Ai_pdm3_F | 92% |
| | | pdm_day2_HW6 | 25% | GT_Ai_pdm3_F and R | GT_Ai_pdm3_F | 84% |

**Table S8. Genotyping results of *pdm3* mKO adult wings.**

| Species | sgRNA | Mutant ID | Wing* | GT primers | Primer used for Sanger sequencing | Indel% |
|---|---|---|---|---|---|---|
| *A. incarnata* | Ai_pdm3_g1 | FM1 | FWR | GT_Ai_pdm3_F and R | GT_Ai_pdm3_F | 90% |
| | | | HWR | GT_Ai_pdm3_F and R | GT_Ai_pdm3_F | 99% |
| | | FW2 | FWR | GT_Ai_pdm3_F and R | GT_Ai_pdm3_F | 46% |
| | | | HWR | GT_Ai_pdm3_F and R | GT_Ai_pdm3_F | 91% |
| | | FW3 | FWR | GT_Ai_pdm3_F and R | GT_Ai_pdm3_F | 97% |
| | | | HWR | GT_Ai_pdm3_F and R | GT_Ai_pdm3_F | 97% |
| | | M1 | FWR | GT_Ai_pdm3_F and R | GT_Ai_pdm3_F | 29% |
| | | | HWR | GT_Ai_pdm3_F and R | GT_Ai_pdm3_F | 62% |
| | | M2 | FWR | GT_Ai_pdm3_F and R | GT_Ai_pdm3_F | 97% |
| | | | HWR | GT_Ai_pdm3_F and R | GT_Ai_pdm3_F | 96% |
| | | M3 | FWR | GT_Ai_pdm3_F and R | GT_Ai_pdm3_F | 96% |
| | | | HWR | GT_Ai_pdm3_F and R | GT_Ai_pdm3_F | 98% |
| *D. plexippus* | Dp_pdm3_g1 | g1_M1 | FWL | GT_Dp_pdm3_F and R | GT_Dp_pdm3_R | 99% |
| | | | HWL | GT_Dp_pdm3_F and R | GT_Dp_pdm3_R | 98% |
| | | g1_FM2 | FWL | GT_Dp_pdm3_F and R | GT_Dp_pdm3_R | 65% |
| | | | HWL | GT_Dp_pdm3_F and R | GT_Dp_pdm3_R | 80% |
| | Dp_pdm3_g2 | g2_M1 | FWR | GT_Dp_pdm3_F2 and R2 | GT_Dp_pdm3_R2 | 86% |
| | | | HWR | GT_Dp_pdm3_F2 and R2 | GT_Dp_pdm3_R2 | fail |
| | | g2_FM2 | FWR | GT_Dp_pdm3_F2 and R2 | GT_Dp_pdm3_R2 | 93% |
| | | | HWR | GT_Dp_pdm3_F2 and R2 | GT_Dp_pdm3_R2 | 95% |
| | | g2_FM3 | FWR | GT_Dp_pdm3_F2 and R2 | GT_Dp_pdm3_R2 | 84% |
| | | | HWR | GT_Dp_pdm3_F2 and R2 | GT_Dp_pdm3_R2 | 92% |

*: Forewing (FW), Hindiwng (HW), Right (R), Left (L) from the ventral view
